# A robust *ex vivo* system to study cellular dynamics underlying mouse peri-implantation development

**DOI:** 10.1101/2021.08.22.457265

**Authors:** Takafumi Ichikawa, Hui Ting Zhang, Laura Panavaite, Anna Erzberger, Dimitri Fabrèges, Rene Snajder, Adrian Wolny, Ekaterina Korotkevich, Nobuko Tsuchida-Straeten, Lars Hufnagel, Anna Kreshuk, Takashi Hiiragi

**Affiliations:** European Molecular Biology Laboratory (EMBL), 69117 Heidelberg, Germany; Institute for the Advanced Study of Human Biology (WPI-ASHBi), Kyoto University, 606-8501 Kyoto, Japan; Collaboration for PhD degree between EMBL and Heidelberg University, Faculty of Biosciences, Heidelberg, Germany.

**Keywords:** Mouse embryonic development, Embryo implantation, Egg cylinder formation, Epiblast morphogenesis, Lumen formation, Tissue-tissue interaction, Mechano-chemical interplay, Embryo culture, *In toto* live-imaging, Quantitative image analysis

## Abstract

Upon implantation, mammalian embryos undergo major morphogenesis and key developmental processes such as body axis specification and gastrulation. However, limited accessibility obscures study of these crucial processes. Here, we develop an *ex vivo* Matrigel-collagen-based culture to recapitulate mouse development from E4.5 to 6.0. Our system not only recapitulates embryonic growth, axis initiation, and overall 3D architecture in 49% of cases, its compatibility with light-sheet microscopy enables study of cellular dynamics through automatic cell segmentation. We find that upon implantation, release of the increasing tension in the polar trophectoderm is necessary for its constriction and invagination. The resulting extra-embryonic ectoderm plays a key role in growth, morphogenesis and patterning of the neighboring epiblast, which subsequently gives rise to all embryonic tissues. This 3D-*ex vivo* system thus offers an unprecedented access to peri-implantation development for *in toto* monitoring, measurement and spatio-temporally controlled perturbation, revealing a mechano-chemical interplay between extra-embryonic and embryonic tissues.

## INTRODUCTION

Implantation is a unique event in mammalian development, whereby an exchange interface is established between the embryo and the maternal tissues (Hemberger et al., 2020; Wang and Dey, 2006). In the first few days following fertilization, the pre-implantation embryo develops into a fluid-filled blastocyst, wherein the pluripotent epiblast is sandwiched between the outer trophectoderm (TE) and the primitive endoderm (PrE). Upon implantation, the extra-embryonic portion of the embryo that consists of TE and PrE-derived cells, engages the maternal tissue in a complex interplay that eventually form the placenta. In the embryo proper, implantation coincides with major changes in tissue architecture as it undergoes gastrulation and body axes specification (Arnold and Robertson, 2009; Rossant and Tam, 2009; Takaoka and Hamada, 2012). While genetic studies characterized key genes and signaling pathways required for these processes, their underlying cellular mechanisms remain obscured by inaccessibility to the implantation process.

*Ex vivo* culture provides an experimental setting to monitor, measure and manipulate embryonic development to glean mechanistic insight. *Ex vivo* culture of peri-implantation mouse embryos so far relied on embryonic growth on 2D surfaces (Bedzhov and Zernicka-Goetz, 2014; HSU, 1971, 1972; Morris et al., 2012; Pienkowski et al., 1974; Tachi, 1992). This culture typically induces adhesion and spreading of trophoblast cells over the surface, disrupting embryonic morphogenesis. Therefore, recapitulation of *in vivo* development is limited with current methods, both in terms of efficiency and physiological relevance.

*In toto* live-imaging has been carried out for later post-implantation mouse development, using light-sheet microscopes (Ichikawa et al., 2013; McDole et al., 2018; Udan et al., 2014; Yue et al., 2020). However, these experimental settings limit the sample number and do not allow quantitative analyses of cellular and tissue morphogenesis or spatio-temporally controlled perturbations. Furthermore, while many studies have introduced new experimental systems focused on the development of embryonic tissues (van den Brink et al., 2014; Deglincerti et al., 2016; Shahbazi et al., 2016; Warmflash et al., 2014; Zheng et al., 2019), the contributions of extra-embryonic or uterine tissues in embryonic development has been relatively overlooked, largely due to technical challenges (Brennan et al., 2001; Christodoulou et al., 2019; Guzman-Ayala et al., 2004; Hiramatsu et al., 2013; Thomas and Beddington, 1996). It remains to be examined how interactions between embryonic and extra-embryonic tissues influence mammalian peri-implantation development.

It is evident that tissue interactions are vital throughout development. Morphogenesis induces spatio-temporally coordinated tissue-tissue interactions that feed back on cellular behaviors such as differentiation and cellular rearrangement, which in turn guides morphogenesis (Bailles et al., 2019; Eiraku et al., 2011; Harland and Gerhart, 1997; Münster et al., 2019; SAUNDERS, 1968; Shyer et al., 2015). To study such an interaction between embryonic and extra-embryonic tissues, it is essential to have an experimental system that faithfully recapitulates peri-implantation development *ex vivo*. Furthermore, the system must be compatible with live monitoring, measurement and perturbation tools to provide mechanistic insights into relevant processes.

In the present study, we developed a 3D culture method for mouse peri-implantation embryos that can couple to *in toto* light-sheet live-imaging. Quantitative analyses of tissue dynamics at single-cell resolution and biophysical measurements and perturbations revealed a key role for mechano-chemical interactions between embryonic and extra-embryonic tissues during early mammalian development.

## RESULTS

### Tension release in the trophectoderm enables invagination and formation of the extra-embryonic ectoderm

With the goal of recapitulating *in utero* development *ex vivo*, we attempted to culture peri-implantation mouse embryos while maintaining their 3D-morphology. Blastocysts developed *in utero* up to embryonic day 4.5 (E4.5) were recovered and embedded into a mixture of Matrigel and collagen. Their development was compared with embryos *in utero* at equivalent stages undergoing implantation and egg cylinder formation. After 24 hours of culture, however, this *ex vivo* condition failed to support proper embryonic development, resulting in disorganized morphology and deterioration of the epiblast (Figure 1A). To improve this culture method, we compared embryos at early stages of culture to corresponding stages of *in utero*-developed embryos. We found that the polar trophectoderm (pTE) cells do not invaginate and thus fail to form the extra-embryonic ectoderm (ExE) after 6 hours in culture, unlike E4.75 embryos developed *in utero* (Figure 1A) (Christodoulou et al., 2019; Copp, 1979). Instead, pTE cells in cultured embryos appeared highly stretched, in contrast to the columnar appearance in embryos developed *in utero* (Figure 1B). This suggested high tension acting on pTE cells in culture. Accordingly, actin and bi-phosphorylated myosin regulatory light chain (ppMRLC) were enriched at the apical surface of the pTE cells before and during invagination in *in utero* developed embryos, whereas they were localized at cell-cell junctions in those developed *ex vivo* (Figures 1C and 1D). Furthermore, direct measurement by micropipette aspiration (Maître et al., 2015) indicated that cortical tension of pTE cells increases during this developmental period (Figure 1E). Collectively, these findings suggest that pTE cells invaginate from the surface layer by the apical constriction, similarly to *Drosophila* gastrulation (Martin et al., 2009), and that excess tension acting on pTE cells, as induced by this culture method, prevents pTE invagination and subsequent ExE formation.

**Figure 1.**
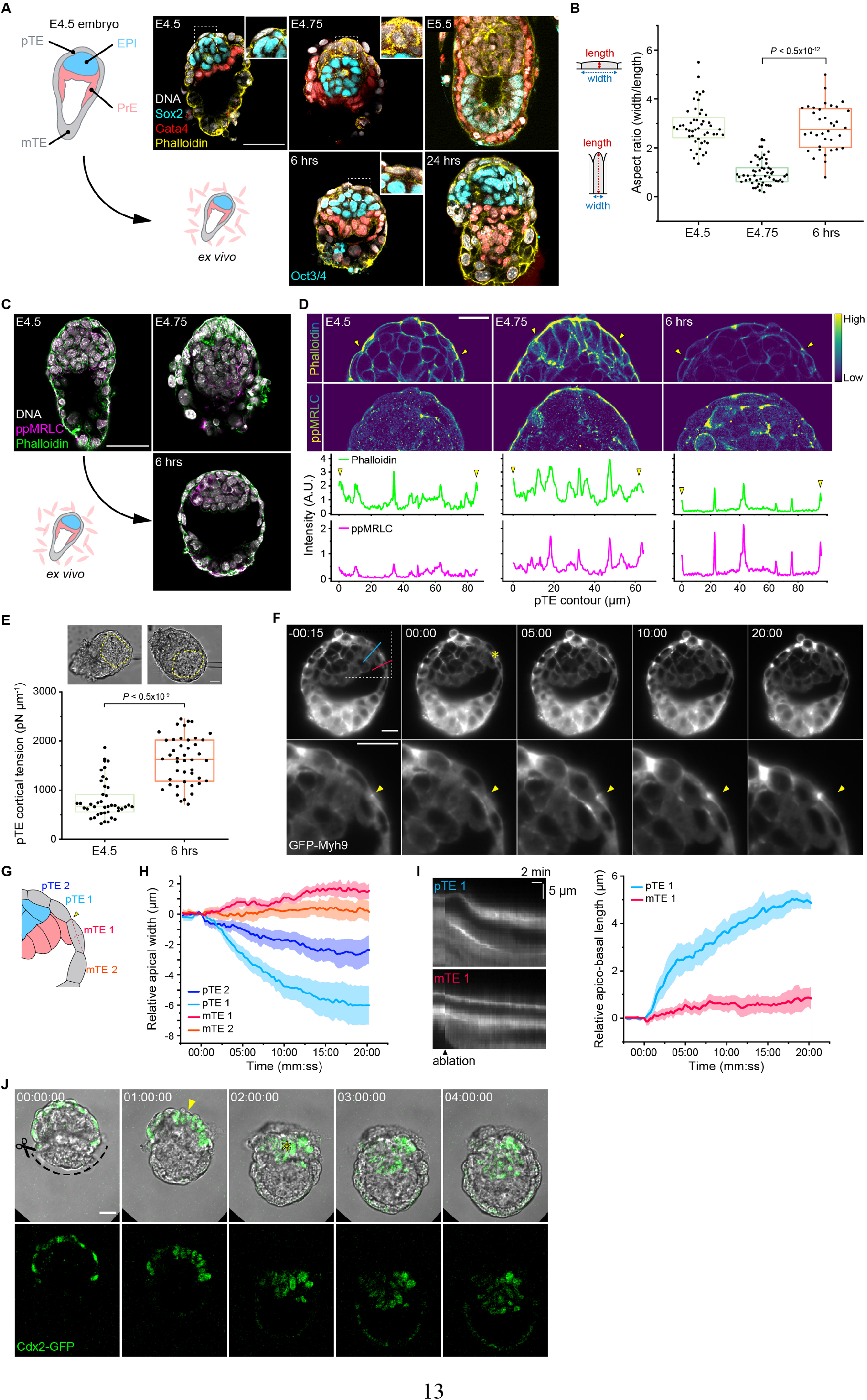
Trophectoderm tension release enables invagination and formation of the extra-embryonic ectoderm. (A) Representative images of mouse embryos developed *in utero* (top) in comparison to those cultured in 3D in Matrigel and collagen mix from E4.5 (bottom) immunostained for Sox2^+^ (E4.5) or Oct3/4^+^ (others) epiblast, Gata4^+^ VE, actin and DNA. *n* = 16 (E4.5), 18 (E4.75), 19 (E5.5), 12 (6hrs) and 18 (24hrs), respectively. (B) Aspect ratio (width to height) of pTE cells in embryos developed *in utero* or in 3D-gel culture shown in (A). *n* = 16, 18 and 12, respectively. Three pTE cells per embryo were measured. (C) Representative images of mouse embryos developed *in utero* (top) in comparison to those cultured in 3D-gel from E4.5 (bottom) immunostained for actin, bi-phosphorylated myosin regulatory light chain (ppMRLC) and DNA. *n* = 24 (E4.5), 34 (E4.75) and 16 (6hrs), respectively. (D) Subcellular localization of actin and ppMRLC along the apical surface of pTE cells in embryos shown in (C). Peaks indicated by arrowheads in the intensity profiles (bottom) correspond to the cell-cell junctions indicated in the microscope images (top). (E) Cortical tension of pTE cells in E4.5 embryos and embryos after 6 hours of culture, measured by micropipette aspiration. *n* = 41 cells from 16 embryos (E4.5) and 44 cells from 18 embryos (6hrs). Yellow broken lines mark the epiblast. (F) Time-lapse images of a representative GFP-Myh9 embryo after 6 hours of 3D-gel culture, ablated with infra-red laser pulses. Ablated point is marked with an asterisk (*t* = 00:00, top) and arrowheads in enlarged views (bottom). Time, minutes:seconds. (G) Schematic for cell shape analysis of TE cells in (F). (H) Change in the apical width of pTE cells, upon laser ablation. Data presented as mean ± s.e.m. *n* = 6. (I) Kymographs of GFP-Myh9 signal along with blue and red lines in (F), and measurement of the apico-basal length of pTE cells upon laser ablation. Data presented as mean ± s.e.m. *n* = 6. (J) Time-lapse images of a representative Cdx2-GFP embryo at E4.5 following microsurgical removal of mTE cells. An arrowhead indicates the apically constricting pTE cells, an asterisk the invaginated ExE cells. Time, hours:minutes:seconds. *n* = 4. *P* values calculated using Kruskal Wallis ANOVA followed by multiple Mann-Whitney *U* test (B) and Mann-Whitney *U* test (E). Scale bars, 50 µm (A and C), 20 µm (D, E, F and J). See also Videos S1 and S2.

This hypothesis predicts that tension release may allow pTE cells to undergo apical constriction and invagination. We tested this by two micromanipulation methods that release TE tension at the boundary between polar and mural TEs, and examined their impact at different spatio-temporal scales. First, spatio-temporally controlled infra-red laser pulses (de Medeiros et al., 2020) targeted at the TE apical cortex of the embryos cultured for 6 hours indeed induced apical constriction of pTE cells over the following 20 minutes (Figure 1F; Video S1). The pTE cells shortened apically and elongated along their apico-basal axis (Figures 1G-I). Next, to examine the impact of tension release at a longer time-scale, we microsurgically excised the mTE from E4.5 embryos at the mTE-pTE boundary. This rapidly induced pTE cell constriction, followed by invagination after 2 hours (Figure 1J; Video S2). These results support that pTE cells invaginate by apical constriction during implantation, which requires release of tension acting on TE cells.

We therefore decided to excise the mTE for subsequent 3D culture of peri-implantation mouse embryos. Notably, this finding that mTE removal improves *ex vivo* culture is in agreement with an earlier observation (Bedzhov et al., 2014), despite the difference between 3D and 2D culture methods. Taken together, these findings show that during mouse peri-implantation development, TE tension increases before its release enables the apical constriction of pTE cells for invagination, growth, and formation of the ExE tissue.

### 3D-geec recapitulates mouse peri-implantation development

Using mTE-removed E4.5 embryos (Figure S1A), we further investigated conditions optimal for robust recapitulation of peri-implantation development *ex vivo* in a 3D gel environment. To this end, we introduced quantitative measures to define the initial embryonic parameters for successful *ex vivo* development and to evaluate the outcome after 48 hours of culture (Figure 2). We found that embryos recovered at E4.5 after natural mating exhibit a high degree of variability in their progression of development, with the combined number of cells in the inner cell mass and pTE ranging from 55 to 232 (Figures S1B and S1C). The success rate of *ex vivo* culture increases with cell number at E4.5. Therefore, the high sample number and efficient *ex vivo* development can be achieved in combination when E4.5 embryos are selected for cell numbers greater than or equal to 110 (Figure S1C). Although this introduces an additional step before culture, in which the embryos are labeled with Hoechst and briefly imaged by confocal microscopy to count cell number, this does not compromise development (Figure S1D). Together, cell number is a reliable predictor of successful *ex vivo* development and ensures consistent and robust experimental outcomes.

**Figure 2.**
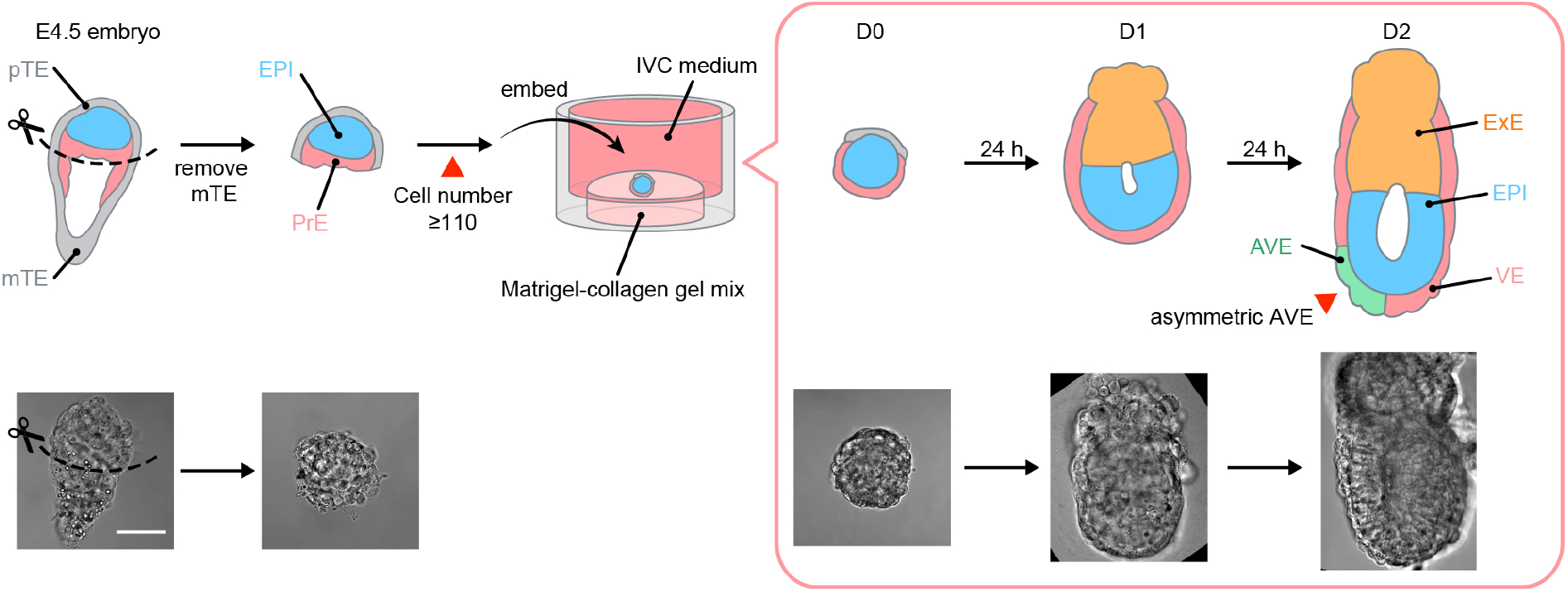
A new 3D *ex vivo* culture of the mouse peri-implantation embryo. Schematic of the 3D-geec workflow (top) and representative DIC images of the embryo (bottom). Mouse embryos from natural mating are recovered at E4.5. After microsurgically removing mTE, the qualified embryos, with cell number 110 or higher (see Figure S1), are embedded in a Matrigel-collagen mix and immersed in IVC medium. Embryo development is evaluated after 48 hours of culture by the formation of egg cylinder (see Figure 3) and the asymmetric distribution of AVE (see Figure 4). Scale bar, 50 µm.

With these embryos we quantitatively evaluated the performance of our 3D *ex vivo* culture, named <u>3D</u>-<u>g</u>el <u>e</u>mbedded <u>e</u>mbryo <u>c</u>ulture (3D-geec), in comparison to embryos developed *in utero*. Over 48 hours, development in 3D-geec closely follows *in utero* developmental changes that occur from E4.5 to E6.0, as judged from embryo morphology and anterior-posterior axis specification (Figures 3A and S2A). Again, we found that *in utero* development progresses with considerable variability in embryo size and cell number (Figures 3B and 3C). The dimensions of 3D-geec embryos are largely comparable, while exhibiting a slightly higher diameter-to-length ratio when compared to *in utero* counterparts (Figure 3D). Cell numbers in the epiblast and visceral endoderm (VE) show a proportional increase in 3D-geec, with 1.5 days (from E4.5 to E6.0) of *in utero* development achieved by 48 hours in culture. Overall, embryonic development in 3D-geec exhibits a 25% temporal delay based on cell number (Figures 3C, 3E and 3F), and these data offer a faithful and quantitative method to stage mouse peri-implantation embryos upon recovery by cell number (Figure S2B; see also Figure 6C).

**Figure 3.**
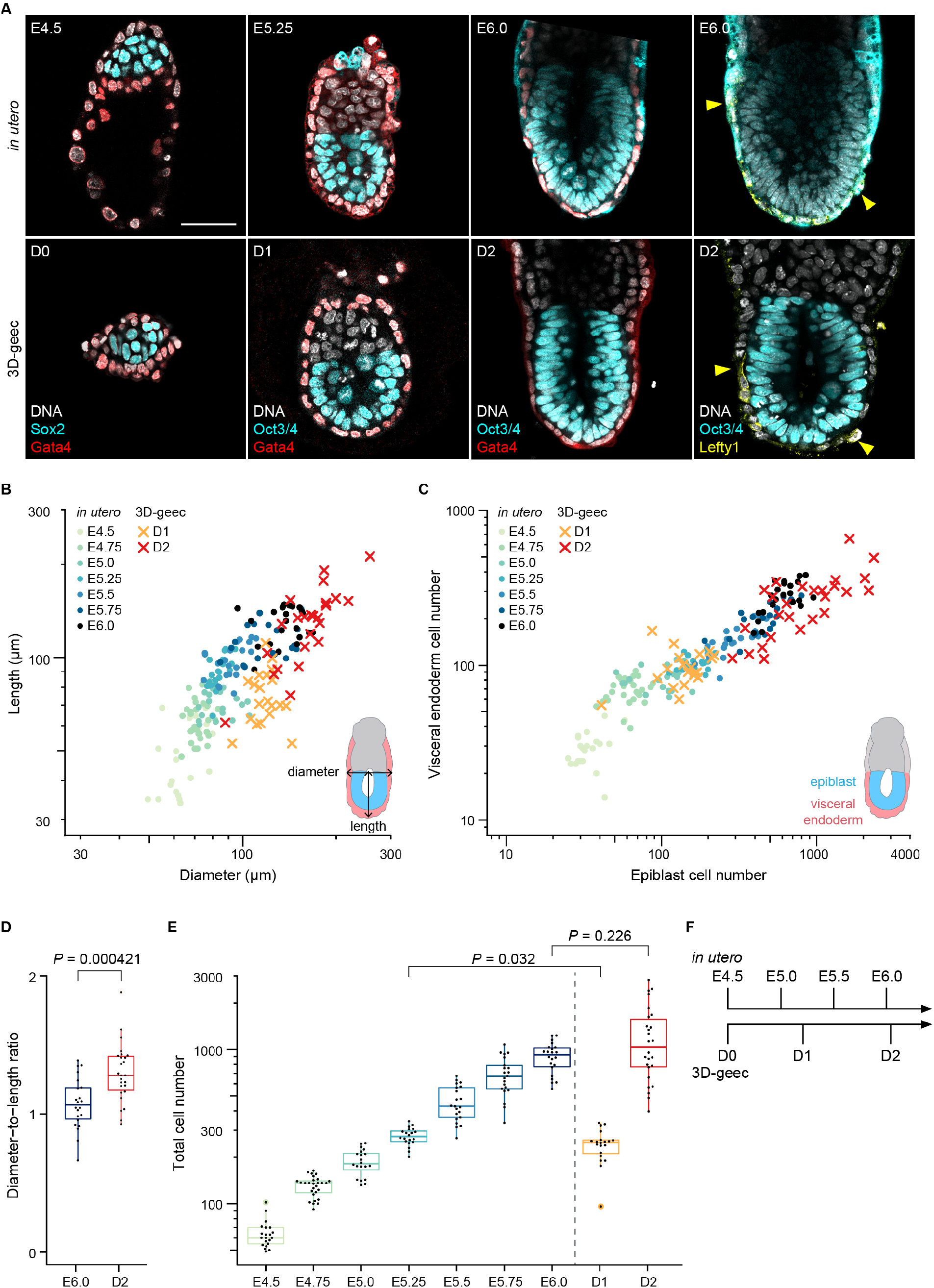
3D-geec recapitulates *in utero* development from E4.5 to E6.0 by 48 hours. (A) Comparison of 3D-geec embryos from day 0 (D0) to 2 (D2; bottom) with embryos developed *in utero* from E4.5 to E6.0 (top). Immunofluorescence images of a representative embryo (see Figure S2 for additional stages) stained for Sox2^+^ or Oct3/4^+^ epiblast, Gata4^+^ VE and asymmetrically localized Lefty1^+^ AVE (yellow arrowheads). (B-C) Scatter plots showing the length against diameter (B), and cell numbers of PrE/VE covering epiblast against epiblast (C) for 3D-geec and *in utero* embryos in log scale. (D) The diameter-to-length ratio for embryos shown in (B). (E) The total (epiblast and PrE/VE) cell numbers for embryos shown in (C). (F) Scaled timeline of 3D-geec development based on the total cell numbers (E). 3D-geec embryos correspond to E5.20 and E6.04 *in utero* embryos after 24 and 48 hours in culture, respectively. *n* = 21 (E4.5), 28 (E4.75), 20 (E5.0), 20 (E5.25), 21 (E5.5), 21 (E5.75), 22 (E6.0), 20 (D1), and 26 (D2). *P* values calculated using *t*-test (D) and Kruskal Wallis ANOVA followed by multiple Mann-Whitney *U* test (E). Scale bars, 50 µm. See also Figure S1 and S2.

We also introduced additional measures to evaluate the outcome of 3D-geec based on cell differentiation and embryonic patterning. For this purpose, we analyzed the distribution of anterior visceral endoderm (AVE) cells in the VE after 48 hours in 3D-geec. At E6.0, mouse embryos establish the anterior-posterior axis by migration of distal visceral endoderm cells to the anterior, forming the AVE (Brennan et al., 2001; Thomas et al., 1998). To quantitatively assess anterior-posterior axis specification, we define an AVE asymmetry index based on the 3D distribution of Lefty1 or Cerl1-expressing AVE cells relative to Gata4-expressing VE cells at the distal tip of the egg cylinder (Figures 4A and 4B). Using *in utero* developed embryos as a reference, we found that 3D-geec embryos with an AVE asymmetry index greater than 0.15 can be considered as having successfully established asymmetric AVE distribution and hence the anterior-posterior embryonic axis (Figure 4C). Based on this criterion, 67% (n=12 of 18) of the 3D-geec-derived egg cylinders display AVE asymmetry. On the other hand, 74% (n=17 of 23) of E4.5 embryos develop to an egg cylinder after 48 hours in 3D-geec. Collectively, these quantitative control measures (see Figure S1; STAR Methods) indicate that 3D-geec recapitulates mouse peri-implantation development to E6.0 with an overall success rate of 49% (Figure 4D).

**Figure 4.**
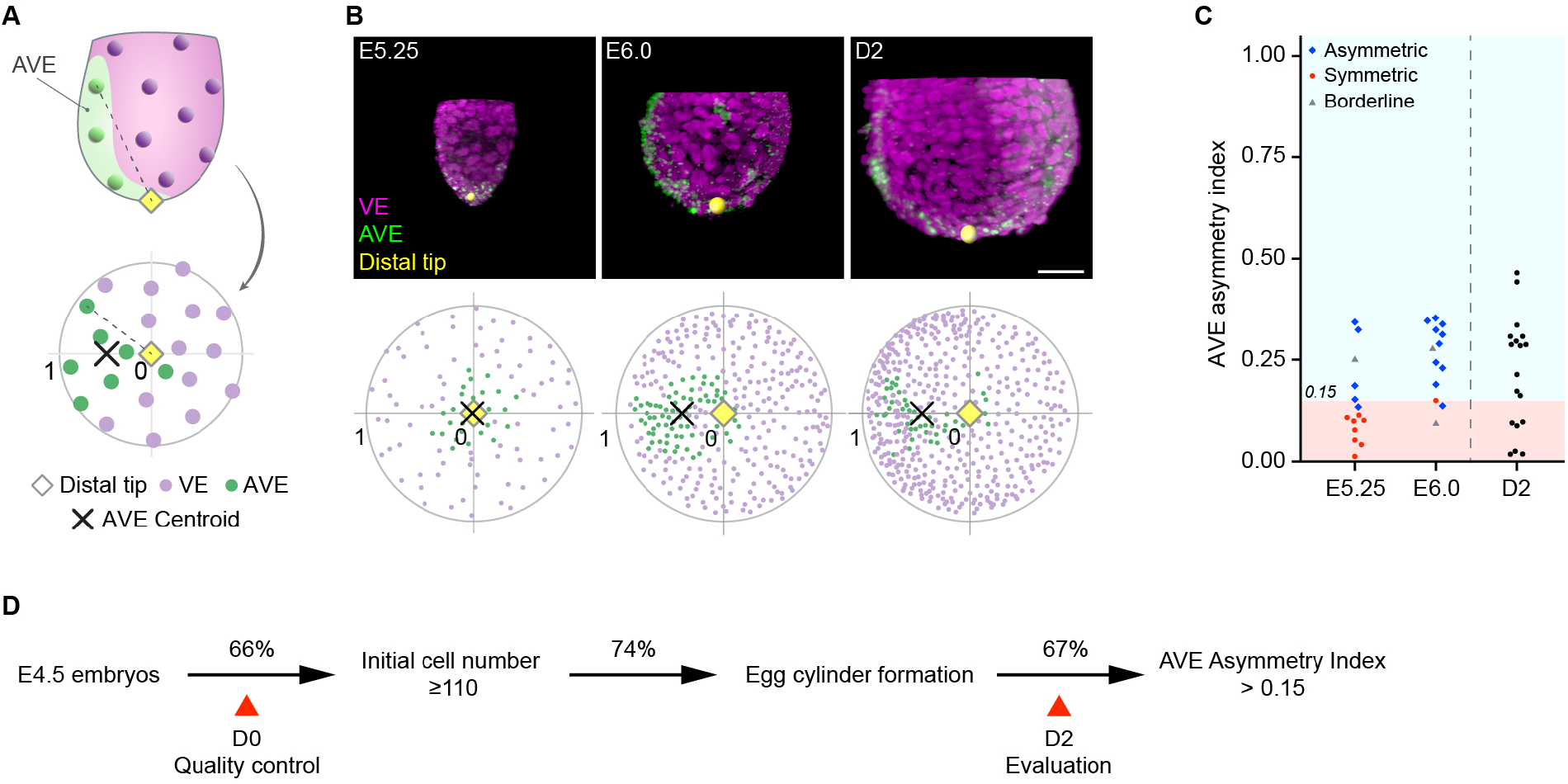
Quantitative evaluation of 3D-geec outcome based on AVE asymmetry at D2. (A) Schematic to calculate the AVE asymmetry index. A polar plot shows the distribution of VE and AVE cells in the embryo, using the distance and angle from the distal tip. The AVE asymmetry index is calculated based on the position of AVE centroid in this polar plot. (B) Representative 3D projections of E5.25, E6.0 and D2 embryos immunostained for Gata4 (VE) and Lefty1 or Cerl1 (AVE), and their respective polar plots. (C) AVE asymmetry index of E5.25, E6.0 and D2 embryos. AVE position of *in utero* embryos was first qualitatively classified as asymmetric (blue), symmetric (red), or borderline (grey). No embryos at E5.25 and E6.0 that were classified as symmetric have an AVE asymmetry index larger than 0.15. Thus, we used this value to evaluate 3D-geec D2 embryos for AVE asymmetry. *n* = 15 (E5.25), 13 (E6.0), and 18 (D2). (D) Summary of 3D-geec efficiency based on quantitative quality control and evaluation. Scale bar, 50 µm. See also Figure S1.

### Live-imaging with an inverted light-sheet microscope reveals cellular dynamics

Next, we aimed to develop *in toto* live-imaging microscopy compatible with 3D-geec. Increasing embryo size, photo-sensitivity, and the long gel-filled distance between the imaging objective and the embryo render our system incompatible with most conventional microscopes. We therefore employed an inverted light-sheet microscope (Strnad et al., 2016) to live-image 3D-geec embryos (Figure 5). The mTE-removed E4.5 embryos were embedded in gel, immersed in IVC medium, and further covered by mineral oil to prevent evaporation (Figure 5A). This setting allows for successful live-imaging of mouse peri-implantation embryos under 3D-geec for 48 hours without compromising embryonic development, as judged from AVE formation, embryo dimensions, and cell number (Figures 5B-F; Videos S3 and S4).

**Figure 5.**
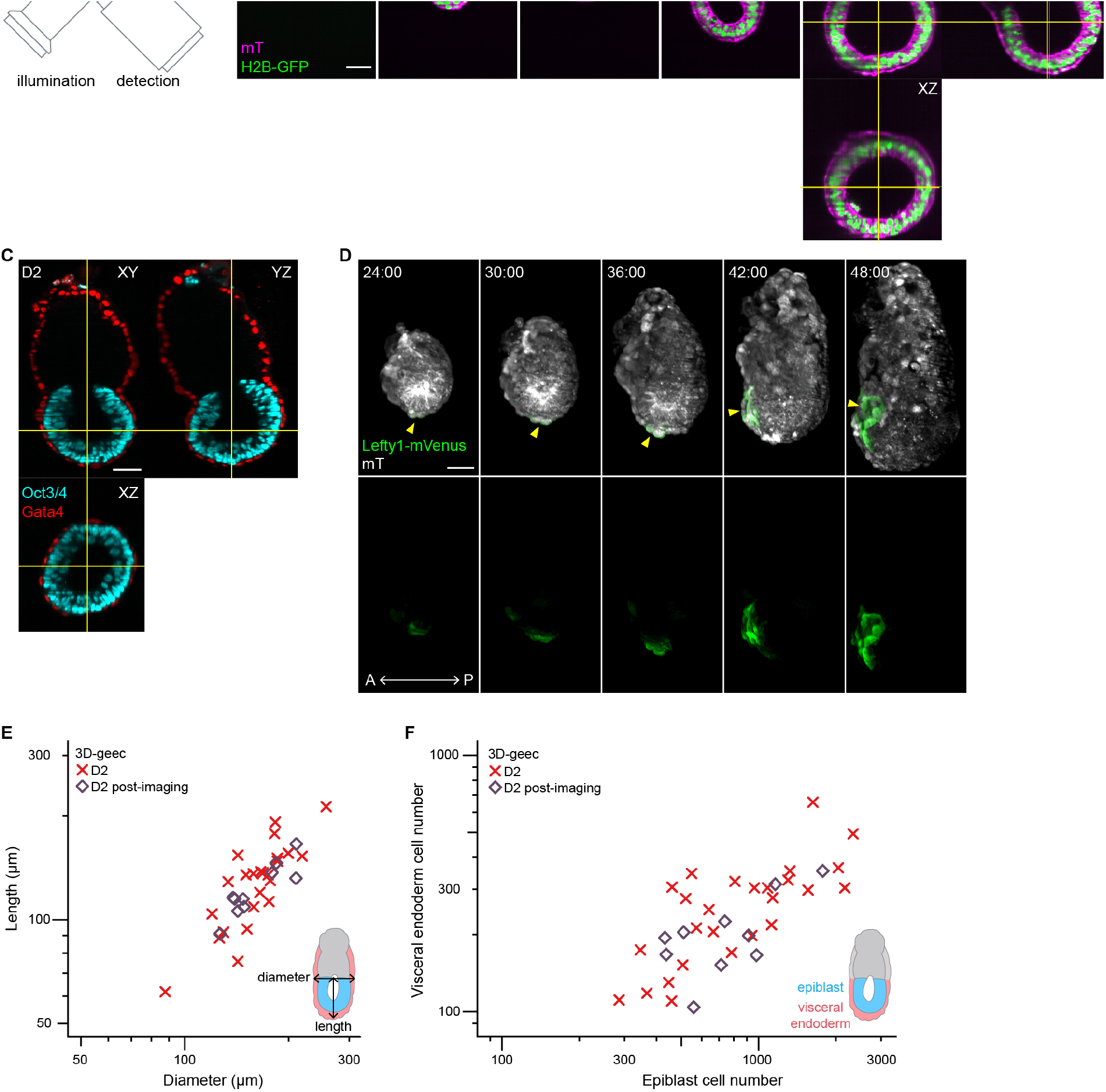
Live-imaging of mouse peri-implantation development during 3D-geec. (A) Design schematic of an inverted light-sheet microscope adapted to 3D-geec. (B) Time-lapse images of a representative H2B-GFP;mT mouse embryo developing from E4.5 (*t* = 00:00, hours:minutes) in 3D-geec with inverted light-sheet microscopy. *n* = 13, *N* = 5. (C) Immunofluorescence of a representative mouse embryo after 48 hours 3D-geec live-imaging (B), stained for Oct3/4^+^ epiblast and Gata4^+^ VE. *n* = 10, *N* = 4. (D) Time-lapse images of a representative Lefty1-mVenus;mT embryo developing in 3D-geec during D1-2. Arrowheads indicate Lefty1^+^ DVE/AVE cells. *n* = 5, *N* = 3. (E-F) Embryo dimension (E) and cell number (F) after 48 hours 3D-geec live-imaging. There is no significant difference between live-imaged embryos (*n* = 10) and embryos cultured in the incubator (*n* = 26, see Figure 3) in terms of diameter (*P* = 0.899), length (*P* = 0.475), epiblast cell number (*P* = 0.689), and VE cell number (*P* = 0.158). *P* values calculated using Mann-Whitney *U* test (diameter and length in (E)), *t*-test (epiblast and VE cell number in (F)). Scale bars, 50 µm. See also Videos S3 and S4.

Quantitative analysis of cellular dynamics is essential for mechanistic understanding of embryonic morphogenesis and patterning. However, available studies so far have limited their analyses to nuclear tracks and lineages (Ichikawa et al., 2013; McDole et al., 2018; Udan et al., 2014; Yue et al., 2020). To extend the analysis to cell shape changes and cell-cell interactions, we developed a machine-learning-based image-processing pipeline for automatic segmentation of cells based on their plasma membrane signal (Figures 6A and 6B; Video S5). We focused on the first 30 hours of 3D-geec culture to quantitatively analyze cellular dynamics of the epiblast tissue. Note that we normalized the developmental stage of individual embryos according to the epiblast cell number at the start and end of live-microscopy (nE; Figure 6C; see Methods, Figure S2B). While cell volume does not exhibit consistent changes during these stages, many epiblast cells undergo elongation, and those elongated cells progressively align radially to form a rosette. The apical domain emerges in epiblast cells and progressively clusters at the center of the epiblast tissue as cells elongate, where the pro-amniotic cavity eventually forms (Figures 6D and 6E; Video S6). Together, quantitative characterization of cellular geometry and polarization reveals their dynamic coordination, suggesting feedback mechanisms underlying maturation of the epiblast, in addition to the key role that epiblast cell polarization plays in pro-amniotic cavity formation (Bedzhov and Zernicka-Goetz, 2014; Christodoulou et al., 2018).

**Figure 6.**
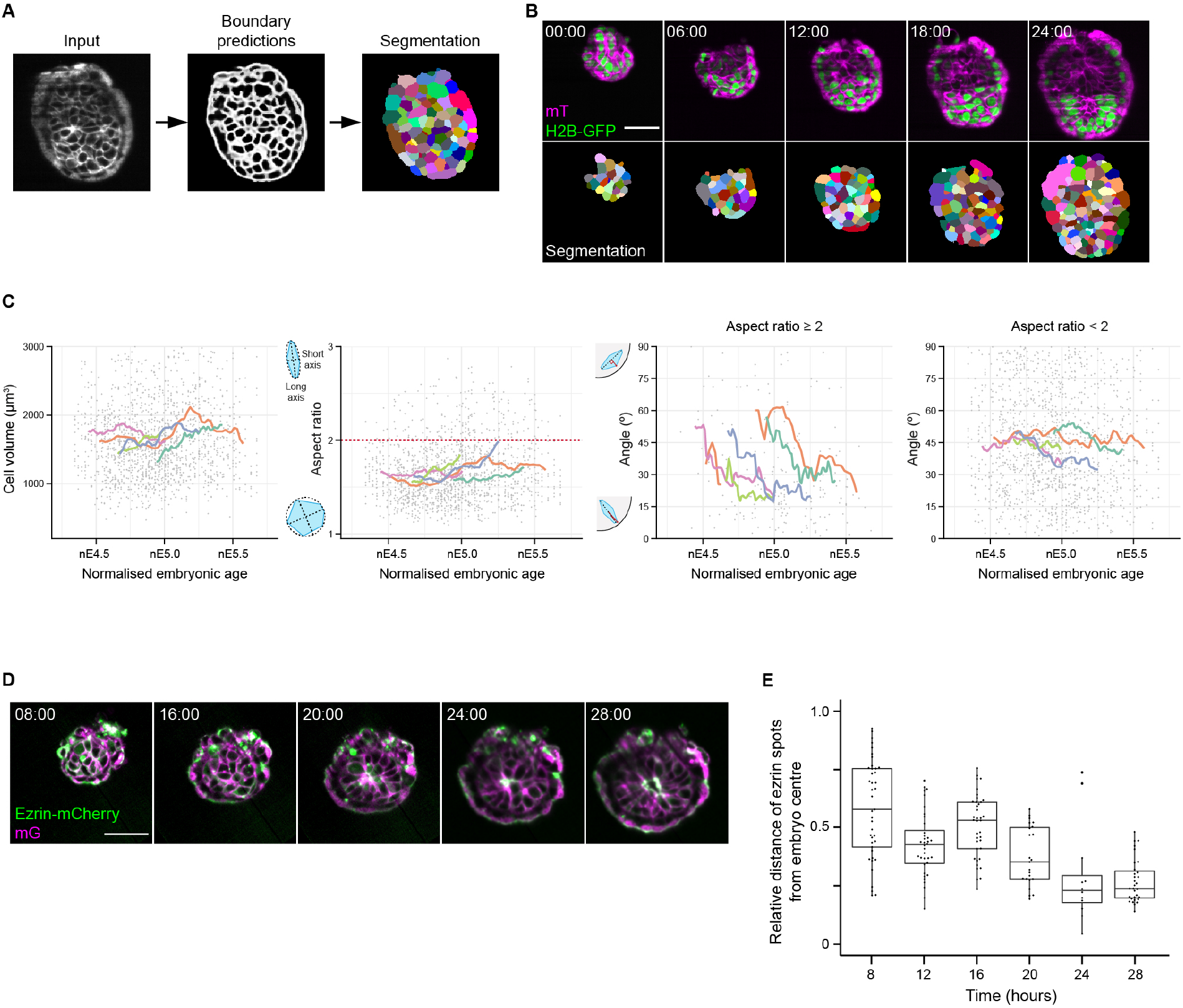
Cellular dynamics underlying mouse peri-implantation morphogenesis and patterning. (A) Schematic explaining the machine-learning-based image-processing pipeline for cell membrane segmentation. (B) Time-lapse images of a representative H2B-GFP;mT mouse embryo developing during the first 24 hours in 3D-geec (top, Figure 5B) and the outcome of automatic cell membrane segmentation (bottom). *n* = 7 (C) Measurement of cell volume, aspect ratio and long-axis alignment against the outer embryonic surface of 5 representative embryos (B) (*n* = 10 cells analyzed for each embryo every hour) for approximately 30 hours of 3D-geec until pro-amniotic cavity formation, with the time-axis normalized according to their epiblast cell numbers (see Figure S2B). (D) Time-lapse images of a representative Ezrin-mCherry;mG mouse embryo developing during the first 28 hours in 3D-geec until pro-amniotic cavity formation. *n* = 8, *N* = 6. (E) Relative distance of the Ezrin signal from the embryo center during 3D-geec (D). Scale bars, 50 µm. See also Videos S5 and S6.

### ExE invagination and growth facilitate epiblast growth, morphogenesis and patterning

While the collective behavior of epiblast cells may thus drive rosette formation, we noted that epiblast cell-cell rearrangement is enhanced upon invagination of pTE cells (Figure S3A). This suggests possible influence by the neighboring ExE tissue on patterning the epiblast. To investigate the impact of ExE on epiblast growth, morphogenesis and patterning, we compared cellular dynamics and morphogenesis between 3D-geec embryos with (see Figure 1) and without mTE (see Figures 2-6). As described earlier, the latter recapitulates ExE formation in a manner comparable to *in utero* development, whereas the former fails to form the ExE, as pTE cells do not invaginate and instead remain a single-layer of cells surrounding the epiblast (Figure 7A; see also Figures 1A at 6 and 24 hours, and 3A at D1). When cell numbers are compared after 18 hours of culture, a properly formed ExE has a higher mean cell number than the pTE that failed to invaginate. Furthermore, the epiblast consists of a significantly higher number of cells in the presence of an ExE (Figure 7B). These data suggest that the presence of the neighboring ExE tissue may facilitate growth of the epiblast.

**Figure 7.**
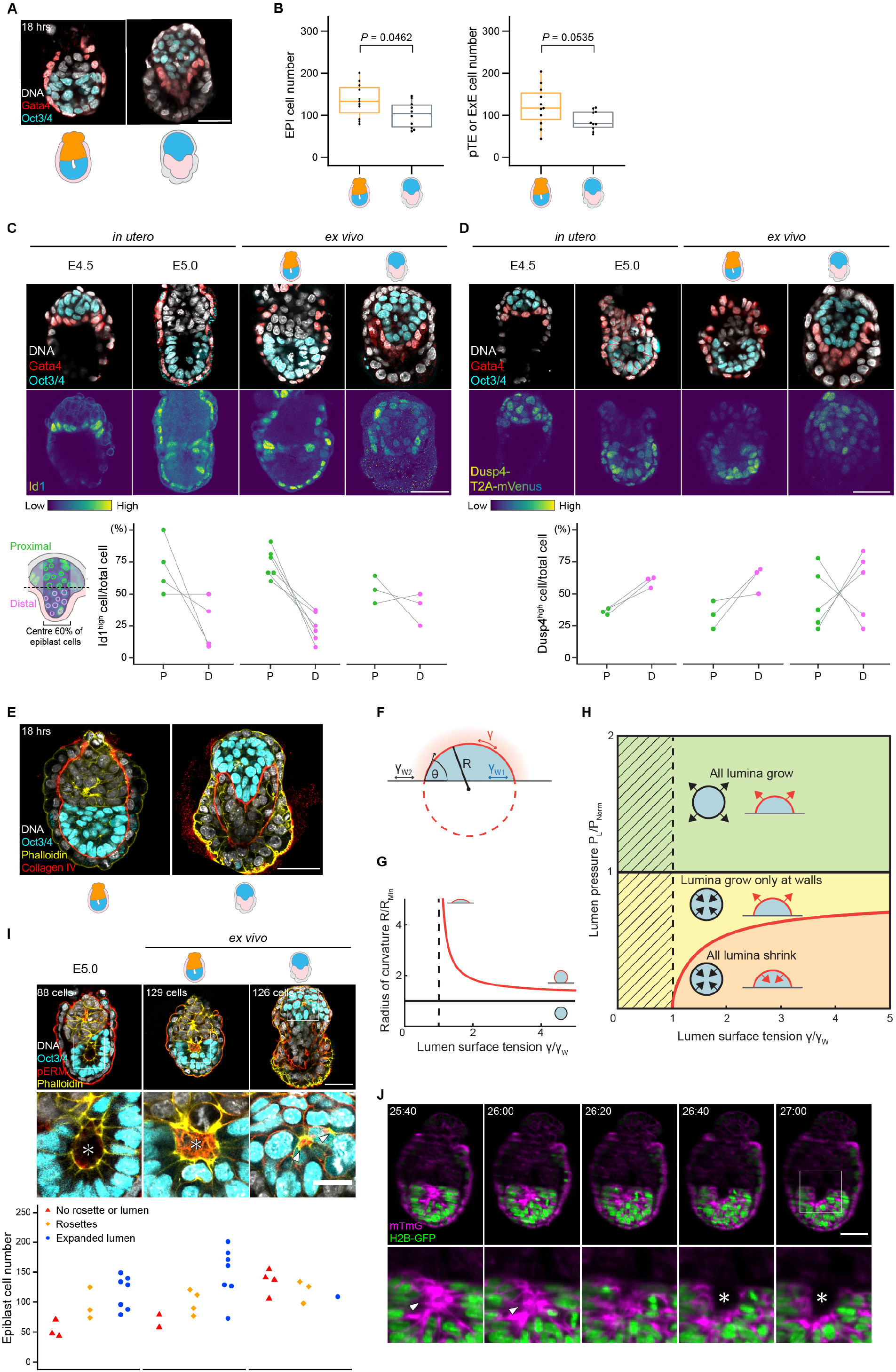
ExE invagination facilitates epiblast growth, morphogenesis and patterning. (A) Immunofluorescence of representative embryos developed *ex vivo* for 18 hours from E4.5 in the presence (left) or absence (right) of ExE stained for Oct3/4^+^ epiblast and Gata4^+^ VE. (B) Epiblast cell numbers (left) and pTE/ExE cell numbers (right) in embryos shown in (A). *n* = 19 and 23, respectively. (C) Immunofluorescence of representative embryos developed *in utero*, or *ex vivo* for 18 hours in the presence or absence of ExE, stained for Id1, Oct3/4^+^ epiblast and Gata4^+^ VE. Percentage of Id1^high^ cells in proximal (P) and distal (D) halves of epiblast of each embryo. *n* = 4, 16, 10 and 14, respectively. (D) Immunofluorescence of representative Dusp4-T2A-mVenus embryos developed *in utero*, or *ex vivo* in the presence or absence of ExE, stained for Oct3/4^+^ epiblast and Gata4^+^ VE. Percentage of Dusp4^high^ cells in proximal (P) and distal (D) halves of epiblast of each embryo. *n* = 4, 5, 6 and 6, respectively. (E) Immunofluorescence of representative embryos developed *ex vivo* for 18 hours from E4.5 in the presence (left) or absence (right) of ExE stained for Collagen IV, actin and Oct3/4^+^ epiblast. (F) Theoretical analysis of lumen formation as a nucleation process. The contact angle θ of a lumen in contact with an external tissue is governed by the surface tensions γ, γW1, γW2 associated with the different interfaces, as described by Young’s equation (see STAR Methods). (G) The radius of curvature of a lumen in contact with a wall is always larger than the radius of curvature of a spherical (homogeneous) lumen of the same volume. RMin refers to the radius of the minimal initial spherical lumen of volume V. The dotted line denotes the point of total wetting. γW = γW2 − γW1. (H) The state diagram shows the parameter regions in which lumina grow or shrink. In the green area, both homogeneous and heterogeneous nucleation events lead to lumen expansion. In the yellow region, only heterogeneous nucleation leads to lumen growth. In the orange region, no lumen growth can be sustained. The hatched region is outside the regime of partial wetting. PNorm = 2γ/RMin. (I) Immunofluorescence of representative embryos developed *in utero*, or *ex vivo* for 18 hours in the presence or absence of ExE, stained for pERM, actin and Oct3/4^+^ epiblast. The number of epiblast cells in relation to the outcome of epiblast patterning (bottom). *n* = 13, 13, and 8, respectively. Arrowheads and asterisks mark the pERM-enriched rosettes and pro-amniotic cavities, respectively. (J) Time-lapse images of a representative H2B-GFP;mT mouse embryo developing *ex vivo* in the presence of ExE from E4.5 (*t* = 00:00, hours:minutes). Arrowheads and asterisks mark rosettes and the pro-amniotic cavity, respectively. *P* values calculated using *t*-test. Scale bars, 50 µm, or 20 µm (enlarged views). See also Figure S3.

To investigate the possible mechanism of this tissue-tissue interaction, we first examined the role of biochemical signaling pathways. Specifically, the expression of Id1, and live-reporter expression of A7-Venus and Dusp4-T2A-mVenus were used to examine BMP, Nodal-Foxh1 (Takaoka et al., 2017), and FGF-Dusp4 signaling pathways, respectively. Nodal signaling in epiblast exhibits cell-to-cell heterogeneity regardless of the presence of ExE (Granier et al., 2011) (Figure S3B). Id1 expression is enriched in the epiblast region in contact with the ExE or pTE, regardless of epiblast geometry or the presence of ExE (Figure 7C), strongly suggesting that BMP is secreted from the ExE or pTE to activate BMP-Id1 signaling in the proximal epiblast, as reported for BMP4 signaling at later stages (Winnier et al., 1995). In marked contrast, FGF-Dusp4 signaling shows an activity gradient across the proximal-distal axis in the epiblast, and this patterned signaling activity is lost in the absence of ExE. This indicates an essential role of ExE in establishing the FGF signaling landscape in the epiblast (Figure 7D). Collectively, these data suggest that biochemical signaling from the ExE acts paracrine to the neighboring epiblast tissue to drive its growth and patterning.

Next, we investigated the possibility of mechanical influence of the ExE tissue on the epiblast. We noted that a flat or convex boundary forms between the ExE and epiblast tissues during pTE invagination and ExE growth (see Figures 3A, 5B, 6B, 7A, 7C, 7D and S3A). This is in contrast to development without ExE, in which the epiblast is surrounded by a single-layer of pTE cells. The boundary between ExE and epiblast lacks Collagen IV, and is therefore distinct from the remainder of the epiblast boundary that is enriched with Collagen IV. The presence of Collagen IV enables Ιntegrinβ1-mediated adhesion at the basal side of epiblast cells (Bedzhov and Zernicka-Goetz, 2014) (Figure 7E). We therefore investigated whether this Collagen-free ExE-epiblast boundary impacts self-organization of the epiblast tissue, particularly the formation of the pro-amniotic cavity.

We started with a theoretical analysis of the mechanics to delineate the conditions under which the cavity can achieve stable growth (STAR Methods). Lumen formation has been described as a process analogous to the nucleation of a droplet in a new phase (Duclut et al., 2019). Briefly, lumina may grow if the inner pressure 𝑃_L_ acting on the lumen-tissue interface is large enough to overcome the resisting interfacial tension:

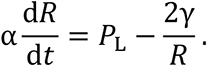

Here 𝑅 denotes the lumen radius, α is a dissipative coefficient, and γ is the effective surface tension associated with the lumen-tissue interface. Notably, the resisting surface term scales inversely with the lumen radius: consequently, the size and shape of a lumen determine its growth. Lumina may grow only if their radius exceeds the critical threshold 𝑅_crit_ = 2γ/𝑃_𝐋_ which is set by the competition of surface and bulk terms. Interestingly, in many physical systems, heterogeneous nucleation – where an external wall or an impurity provides an additional interface for newly forming droplets – dominates over homogeneous nucleation. It is an everyday observation that gas bubbles first form at the surface of a pot of water about to boil. Indeed, the radius of curvature of a nascent droplet adhered to a surface is larger compared to a non-adherent droplet of equal volume (Figures 7F-G). Correspondingly, the surface term resisting nucleation is lowered for an adherent droplet (Turnbull, 1950). We hypothesized that the neighboring ExE tissue provides an interface with properties facilitating lumen formation, analogous to heterogeneous nucleation. We calculated the parameter regimes in which stable lumen expansion is achieved, and found that our theory predicts more robust lumen formation in the presence of the ExE-epiblast boundary (Figure 7H).

To test this prediction, we examined the formation of the pro-amniotic cavity in the embryo developing in the presence or absence of the neighboring ExE tissue. More than half (7 of 13) of embryos developing with an ExE formed a pro-amniotic cavity after 18hrs of 3D-geec, comparable with the 7 of 13 *in utero*-developed E5.0 embryos that had a pro-amniotic cavity (Figure 7I). In contrast, the epiblast is disorganized in embryos that failed to form an ExE, marked by multiple rosettes without a lumen. Regarding the positioning of the nascent pro-amniotic cavity, 38 (90%) of 42 embryos developed in 3D-geec formed the pro-amniotic cavity at the boundary between epiblast and ExE tissues (see Figures 5B, 6B and 7I). Of 36 embryos developed *in utero* (those with a nascent lumen at E4.75, 5.0 and 5.25), 24 (67%) formed the pro-amniotic lumen at the boundary between epiblast and ExE tissues (Figure S3C), with an additional 4 embryos that had already begun epiblast elongation with the lumen connected to the boundary as a fissure-like structure (Figure 7I; a total of 28 embryos, 78%). Furthermore, time-lapse images of 3D-geec embryos show that the lumen emerges at the epiblast-ExE boundary (Figure 7J), or alternatively, that while rosettes form at multiple locations within epiblast tissue, the lumen expands from the epiblast-ExE boundary (Figures S3D and S3E). These data are all in agreement with our calculated prediction that lumen formation is more stable at the epiblast-ExE tissue boundary, illustrating a mechanical contribution of the ExE in shaping the epiblast.

Taken together, these findings strongly suggest that the ExE acts both biochemically and mechanically on the neighboring epiblast to facilitate its growth, morphogenesis and patterning, leading to epiblast elongation and ultimately egg cylinder formation.

## DISCUSSION

The new 3D-geec methods presented here enable 3D-culture and live-imaging of mouse peri-implantation embryos from E4.5 to E6.0 *ex vivo* over 48 hours. Quantitative measures ensure efficient success rate and reproducibility. Our findings show that 3D gel culture recapitulates *in utero* development more faithfully than available 2D methods, in line with a recent study that developed human peri-implantation embryo culture in 3D (Xiang et al., 2020). These 3D gel cultures effectively prevent disruption of embryonic morphogenesis, which has been largely inevitable in the 2D culture methods. Our 3D culture method is also compatible with live-imaging using commercially available light-sheet microscopes (Serra et al., 2019; Strnad et al., 2016), offering sufficient spatio-temporal resolution for automatic image analyses including cell tracking and membrane segmentation. Notably, these microscopes also accommodate multi-sample imaging and enable spatio-temporally controlled perturbations such as photo-manipulation. The unprecedented access to cellular dynamics during peri-implantation development and the capability for *in toto* monitoring, measurement and manipulation will ultimately lead to finer mechanistic understanding of this crucial period in mammalian development.

In this study, we found that releasing TE tension enables pTE cells to invaginate by apical constriction and proliferate to form the ExE. This ExE in turn facilitates the growth and morphogenesis of the neighboring epiblast by establishing BMP and FGF signaling landscapes, consistent with the essential role of BMP signaling in maintaining epiblast pluripotency (Di-Gregorio et al., 2007). The reduced growth of epiblast in the absence of ExE might also be explained by the role of proprotein convertases, Spc1 and Spc4 secreted by ExE, in activating Nodal signaling in the epiblast (Beck et al., 2002). Furthermore, theoretical analysis and biological experiments consistently show that juxta-positioning of the ExE tissue also facilitates luminogenesis and patterned morphogenesis of the epiblast tissue. This is in line with the findings that while embryonic stem cells (Bedzhov and Zernicka-Goetz, 2014) and embryos lacking an ExE (Figures 1 and 7) may form rosettes or a pro-amniotic cavity through intrinsic properties of epiblast cells, epiblast patterning occurs more consistently and robustly *in vivo* through tissue-tissue interactions.

Overall, our data demonstrate biochemical and mechanical roles of extra-embryonic tissues in mammalian embryonic development, in addition to their impact on visceral endoderm morphogenesis, which was reported in a recent study (Christodoulou et al., 2019). As the epiblast is known to control the proliferation of the ExE via FGF (Christodoulou et al., 2019; Gardner et al., 1973), our findings suggest that reciprocal interactions between embryonic and extra-embryonic tissues (Brennan et al., 2001) start as early as E5.0.

While 3D-geec offers novel access to mouse peri-implantation development, it also leaves us with new challenges. First, the present methods require removal of the mTE, similarly to 2D methods (Bedzhov et al., 2014), to release TE tension and induce formation of the ExE. While this results in robust *ex vivo* recapitulation of mouse peri-implantation development from E4.5 to E6.0 in 3D-gel, *in utero* development certainly involves the mTE, which together with the parietal endoderm forms Reichert’s membrane (Salamat et al., 1995). The exact mechanism of how this works *in utero*, and development of an *ex vivo* culture method that retains the mTE, will be topics of future studies. Second, the diameter of the 3D-geec embryo is wider than embryos developed *in utero* (Figure 3B). This spherical tissue dimension may be driven by pressurized expansion of the pro-amniotic cavity, similar to blastocyst cavity expansion prior to implantation (Chan et al., 2019; Dumortier et al., 2019; Leonavicius et al., 2018; Niwayama et al., 2019). Moreover, uterine tissue may be necessary to confine this expansion for precise epiblast elongation during *in utero* development.

In both cases, it is highly conceivable that mammalian peri-implantation development requires intimate interactions between embryos and extra-embryonic tissues, as well as contribution by uterine tissues. Further development of methods to recapitulate these interactions *ex vivo*, or to study embryonic development *in situ* inside the uterus (Huang et al., 2020), will be necessary to integrate their respective roles and gain a comprehensive understanding of mammalian development.

## Supplemental Information

Supplemantal Information includes three figures, one table and six videos.

## Acknowledgements

We are grateful to the members of the Hiiragi lab for discussions and comments on the manuscript, in particular Prachiti Moghe and Chii Jou Chan for instructing cortical tension measurement and Ramona Bloehs and Stefanie Friese for their technical support, and the EMBL animal facility for their support. We thank Jérôme Collignon, Aissam Ikmi and Tristan Rodriguez for critical reading of the manuscript, Hiroshi Hamada and Katsuyoshi Takaoka for Lefty1-mVenus and A7-Venus mouse lines, Robert S. Adelstein for GFP-Myh9 mouse line, Daniel Messerschmidt for peri-implantation embryo recovery expertise, and Luxendo for their technical support in light-sheet microscopy and laser ablation. T.I. is supported by JSPS Overseas Research Fellowship, HT.Z. by the A*STAR National Science Scholarship (NSS PhD) and D.F. by fellowships from the EMBL Interdisciplinary Postdoc Program (EIPOD) under Marie Sklodowska-Curie Actions (COFUND III RTD). The Hiiragi laboratory is supported by EMBL and the European Research Council (ERC Advanced Grant “SelforganisingEmbryo”, grant agreement 742732).

## Author Contributions

T.I., HT.Z. and T.H. designed the study; T.I. and HT.Z. conducted the experiments, and analyzed and interpreted the data together with T.H.; L.P. initiated the project and developed an earlier version of the culture methods; A.E. conceived and conducted theoretical analysis; D.F. developed the computational image analysis together with R.S., A.W. and A.K.; E.K. and N.T.-S. generated transgenic mice; L.H. helped with light-sheet microscopy and laser ablation; T.I. and HT.Z. prepared figures, tables, movies and methods, and T.H. wrote the manuscript with inputs from all authors.

## Supplemental Figures

## STAR★METHODS

### RESOURCE AVAILABILITY

#### Lead Contact

Further information and requests for resources and reagents should be directed to and will be fulfilled by the Lead Contact, Takashi Hiiragi.

#### Materials Availability

All unique/stable reagents generated in this study are available from the Lead Contact with a completed Materials Transfer Agreement.

#### SData and Code Availability

All datasets/codes generated during this study are available upon request.

### EXPERIMENTAL MODEL AND SUBJECT DETAILS

#### Animal Work

All animal work was performed in the Laboratory Animal Resources (LAR) at the European Molecular Biology Laboratory (EMBL) with permission from the Institutional Animal Care and Use Committee (IACUC) overseeing the operation (IACUC number TH11 00 11). LAR is operated according to the Federation of European Laboratory Animal Science Associations (FELASA) guidelines and recommendations. All mice were maintained in specific pathogen-free conditions with 12-12 hours light-dark cycle and used for experiments at the age of 8 to 35 weeks.

#### Mouse Lines and Genotyping

The following mouse lines were used in this study: a F1 hybrid strain between C57BL/6 and C3H (B6C3F1) as wild-type (WT), mTmG (Muzumdar et al., 2007), H2B-GFP (Hadjantonakis and Papaioannou, 2004), GFP-Myh9 (Zhang et al., 2012), Cdx2-GFP (McDole and Zheng, 2012), Lefty1-mVenus (Takaoka et al., 2011), and A7-Venus (Takaoka et al., 2017). Ezrin-mCherry and Dusp4-T2A-3xmVenus were generated in this study. Standard tail genotyping procedures were used to genotype transgenic mice (for primers and PCR product sizes, see Table S1).

To generate Ezrin-mCherry mice, human ezrin coding sequence tagged with mCherry was PCR amplified using pRN3-Ezrin-mCherry plasmid (a gift from Sophie Louvet-Vallée; (Dard et al., 2001)) as a template and primers introducing Nhe1 and EcoR1 recognition sites at the ends of the amplicon. The PCR fragment was digested with Nhe1 and EcoR1 and then inserted into pgk-ATG-FRT2-CAG plasmid (a gift from Judith Reichmann, Ellenberg lab in EMBL) cut with the same restriction enzymes. The obtained plasmid contained Ezrin tagged with mCherry under CAG promoter. This plasmid was digested with Pvu1, and the resulting 7018 bp fragment was used for pronuclear injection into C57BL/6 zygotes to generate a mouse with random integration.

To generate Dusp4-T2A knock-in alleles, we targeted the stop codon of endogenous Dusp4 locus with one selection and two different reporter cassettes coding for a destabilized triple mCherry and triple mVenus. The reporter cassettes were flanked by loxP- and FRT-sites to remove the selection cassette. Thus, the targeting vector was constructed as follows: *loxP-T2A-3xmCherry-NLS-PEST-FRT-PGK Neo-loxP-T2A-3xmVenus-NLS-PEST-FRT*. Cre-mediated excision resulted in Dusp4-T2A-3xmVenus allele used in this study. Dusp4-T2A-3xmVenus knock-in reporter line was generated by standard gene targeting techniques using R1 embryonic stem cells. Briefly, chimeric mice were obtained by C57BL/6 blastocyst injection and then outbred to establish the line through germline transmission.

#### Mouse Embryos

To obtain mouse peri- or early post-implantation embryos, mice were naturally mated, and noon on the day when a vaginal plug was detected was defined as embryonic day 0.5 (E0.5). All embryos were recovered from dissected uteri in dissection medium (DMEM (Gibco, 11880028) supplemented with 15% heat-inactivated FBS (PAA, A15-080), 2 mM GlutaMAX (Gibco, 35050061), 10 mM HEPES (Sigma, H0887), 25 units/mL Penicillin and 25 µg/mL Streptomycin (Gibco, 15070063)) under a stereomicroscope (Zeiss, StreREO Discovery.V8) equipped with a thermo plate (Tokai Hit) at 37°C, as described (Nagy et al., 2003). The Reichert’s membrane of the post-implantation embryos was removed using sharp needles (BD eclipse, 305757). Recovered embryos were handled using an aspirator tube (Sigma, A5177) equipped with a glass pipette pulled from glass micropipettes (Blaubrand intraMark 708744).

### METHOD DETAILS

#### Embryo Culture

Mouse embryos were cultured in an incubator with a humidified atmosphere of 5% CO2 at 37 °C (Thermo Scientific, Heracell 240i). Gel mix for embedding was prepared on ice, first adding basal medium (advanced DMEM/F-12 (Gibco, 12634010) supplemented with 2 mM GlutaMAX, 25 units/mL Penicillin and 25 µg/mL Streptomycin), growth factor reduced Matrigel (Corning, 356230, lot. 7345012), and then rat-tail Collagen I (Corning, 354236, lot. 6053001). Due to lot-to-lot variation of Matrigel and Collagen I, it is recommended to test for culture side-by-side prior to a large purchase. We tested three lots of Matrigel (lot. 7107329, 7202001, and 7543012) and two lots of Collagen I (lot. 5064009 and 6053001) and selected as above based on the rate of successful egg cylinder formation. We also examined different combinations of the final concentration of Matrigel and Collagen I ranging from 0.5 to 5.0 mg/mL and 0 to 0.7 mg/mL, and found 3.0 mg/mL and 0.3 mg/mL resulted in the best performance, respectively. 15 µL gel mix was added in an inner well of the µ-Slide Angiogenesis dish (Ibidi, 81506), and then embryos quickly rinsed with the gel mix were carefully embedded in the gel droplet so that they did neither adhere to the surface of the dish nor float at the interface of the gel. After solidification of the gel upon 30 minutes incubation in the incubator, 50 µL pre-warmed IVC1 medium (Bedzhov et al., 2014) was added to fill the upper well. IVC1 medium was exchanged for IVC2 medium (Bedzhov et al., 2014) after 24 hours of culture.

For 3D gel-embedded embryo culture (3D-geec), mural trophectoderm (mTE) was microsurgically removed from E4.5 embryos immediately after recovery using sharp needles under a stereomicroscope. Then mTE-removed embryos were embedded as described above. To count the initial number of cells, mTE-removed embryos were incubated in IVC1 containing 5 µg/mL Hoechst 33342 (Invitrogen, H21492) for 30 minutes at 37^°^C. Embryos were rinsed with IVC1 three times and live-imaged with 405 nm laser on a confocal microscope (Zeiss, LSM880) in a custom-made incubation box set to 5% CO2 and 5% O2 at 37 °C. To minimise the UV-damage, imaging was achieved within 30 seconds by using Airyscan Fast mode. This additional step before culture ensures the highest quality and consistent experimental outcome without compromising the development (Figure S1D).

#### Cortical Tension Measurement

Micropipette aspiration set-up was used as described previously (Biro and Maître, 2015; Maître et al., 2015) to measure the cortical tension of pTE cells. Briefly, microforged micropipettes coated with Sigmacote (Sigma, SL2) of radius 3-4 µm were coupled to a microfluidic pump (Fluigent, MFCS-VAC). Pressures were increasingly applied in a step-wise manner, until reaching a cortex deformation which has the radius of the micropipette in use (*R*p). At steady state, the cortical tension γ of the pTE is calculated based on Young– Laplace’s law: γ = *P*c/2(1/*R*p − 1/*R*c), where *P*c is the pressure used to deform the cell of radius *R*c. Embryos were cultured in suspension by hanging-drop of IVC1 medium for 6 hrs, prior to micropipette aspiration. The surface of the glass-bottom dish was also coated with Sigmacote to prevent the embryos from attachment to the dish. Microscopic inspection of cell membrane deformation ensured aspiration of a single pTE cell.

#### Immunofluorescence Staining and Imaging

Embryos were fixed with 4% paraformaldehyde (Electron microscopy sciences 19208) in PBS for 15 minutes (*in utero* developed embryos) or 30 minutes (*ex vivo* cultured embryos) at room temperature and subsequently permeabilized with 0.5% Triton X-100 (Sigma, T8787) in PBS for 30 minutes at room temperature with gentle agitation. Embryos were incubated in blocking buffer (5% donkey serum (Sigma, D9663), 2.5% BSA (Sigma, A9647), 0.05% Triton X-100 in PBS) overnight at 4^°^C with gentle agitation. Embryos were then incubated with primary antibodies diluted in the blocking buffer overnight at 4^°^C or 2 hours at room temperature. After washing with the blocking buffer, embryos were further incubated with secondary antibodies diluted in the blocking buffer for 2 hours at room temperature. Dye staining was simultaneously performed with the secondary antibody staining, using DAPI (Invitrogen, D3571) at 10 µg/mL or Rhodamine Phalloidin (Invitrogen, R415) diluted at 1:400. Finally, stained embryos were mounted in PBS.

Primary antibodies against Oct3/4 (Santa Cruz Biotechnology, sc-5279), Gata4 biotinylated (R&D systems, AF2606), Sox2 (Cell Signaling, 23064), and Cdx2 (Biogenex Laboratories, MU392AUC), and Collagen IV (Millipore, AB756P) were diluted at 1:200. Primary antibodies against di-phosphorylated myosin regulatory light chain (ppMRLC) (Cell Signaling, 3674), and phosphorylated ERM (pERM) (Cell Signaling, 3726) were diluted at 1:100. Primary antibodies against Lefty (R&D systems, AF746), Cerberus1 (R&D systems, MAB1986), and Id1 (Biocheck, BCH-1/195-14) were diluted at 1:50.

Secondary antibodies, donkey anti-goat IgG Alexa Fluor 488 (Invitrogen, A11055), donkey anti-goat IgG Alexa Fluor Plus 680 (Invitrogen, A32860), donkey anti-rabbit IgG Alexa Fluor Plus 488 (Invitrogen, A32790), donkey anti-mouse IgG Alexa Fluor Plus 594 (Invitrogen, A32744), donkey anti-mouse IgG Cy5 AffiniPure (Jackson ImmunoResearch, 715-175-150), donkey anti-rat IgG Cy5 AffiniPure (Jackson ImmunoResearch, 712-175-153) were used at 1:200.

Images of immunostained embryos were obtained by LSM880 equipped with a C-Apochromat 40x/1.2 NA water immersion objective (Zeiss). ASE-YFP and Dusp4-mVenus signals were imaged by LSM confocal mode. Otherwise, Airyscan Fast mode was used, and raw Airyscan images were post-processed by ZEN black software (Zeiss). E6.0 and D2 embryos in Figure 4B, and embryos after 48 hours of live-imaging in Figure 5C were imaged by an inverted light-sheet microscope (Bruker, Luxendo, InVi SPIM) to illuminate in deep.

#### Confocal Live-imaging

After removal of mTE, Cdx2-GFP embryos were mounted in 10 µL IVC1 drops covered with mineral oil (Sigma, M8410) on 35 mm glass-bottom dishes (MatTek, P35G-1.5-14-C). Live-imaging was performed on a confocal microscope (Zeiss, LSM780) equipped with a custom-made incubation box set to 5% CO2 and 5% O2 at 37 °C, and a C-Apochromat 40x/1.2 NA water immersion objective. Images were acquired every 5 minutes with 13 Z-slices separated by 5 µm.

#### Light-sheet Live-imaging

3D-geec embryos were live-imaged using InVi SPIM. Up to ten embryos were embedded in a 10 µL gel mix within the V-shaped sample holder covered with transparent FEP foil, carefully positioned so that they are at proximity but do not attach to the FEP foil which would disrupt morphogenesis via adhesion. After gelification, embryos were immersed in 75 µL IVC1 medium and further covered with 200 µL mineral oil to prevent evaporation. IVC1 medium was exchanged for IVC 2 medium after 24 hours of culture. The sample holder was enclosed in an environmentally controlled incubation box with 5% CO2 and 5% O2 at 37 °C.

InVi SPIM was equipped with a Nikon 25x/1.1NA water immersion detective objective and a Nikon 10x/0.3 NA water immersion illumination objective. The illumination plane and focal plane were aligned before each imaging session and maintained during the imaging. Images were taken every 20 min by a CMOS camera (Hamamatsu, ORCA Flash4.0 V2) with line-scan mode in LuxControl (Luxendo). The imaged volume in case of 48 hours of continuous live-imaging was 425.98×425.98×400 µm^3^, with a physical voxel size of 0.208×0.208×1.000 µm^3^, along the X, Y and Z axis, respectively. For the live-imaging shorter than 24 hours, the volume was 212.99×212.99×200 µm^3^ with a physical voxel size of 0.104×0.104×1.000 µm^3^. The lasers and filters used were 488 nm and BP525/50, 515 nm and BP545/40, 561 nm and LP561, and 594 nm and BP632/60 to image GFP, mVenus, tdTomato, and mCherry fluorophores, respectively. Exposure time for each plane was set to 50 ms.

Eighty-one % (n=13 of 16) embryos that expressed both H2B-GFP and mT, alternatively 77% (n=23 of 30) embryos that expressed mT regardless of H2B-GFP, developed into the egg cylinder after 48 hours of live-imaging from 5 independent experiments (Figure 5B) without substantial change in embryo size and cell number (Figures 5E and 5F), suggesting no significant harmful effects of live-imaging on 3D-geec development.

#### Laser Ablation with Light-sheet Live-imaging

To perform laser ablation during light-sheet live-imaging, we equipped InVi SPIM with a photomanipulation module (Bruker, Luxendo) (de Medeiros et al., 2020). Specifically, a pulsed infrared (IR) laser at 1040 nm, 200 femtoseconds pulse length and 1.5W (Spectra-Physics, HighQ-2) was coupled with the detection objective. The illumination spot of IR laser was aligned at the focal plane before each experimental session and maintained during the experiment to ensure spatial control of the ablation while avoiding wound response. Viability of ablated embryos was verified by embryo growth at 6 hours of culture after ablation.

Ablation at the cell-cell junction in a TE layer was performed by defining a circular ROI of 0.8 µm in diameter on the GFP-Myh9 enriched cell-cell junction, and using 100% laser power, 100 ms dwell time, 5 times repetitions and 2 pixels spacing in LuxControl.

Images were taken every 15 seconds with 5 Z-slices separated by 1 µm. Only those experiments in which laser ablation did not elicit typical wound responses such as cortex blebbing, cell swelling or bursting were considered for analysis.

#### Nucleation Theory of Lumen Formation

Fluid-filled cavities can appear within cellular assemblies, and such lumina play important roles during embryonic development (Ryan et al., 2019). Lumen formation has been described as a process analogous to the nucleation of a droplet in a new phase (Duclut et al., 2019), where the competition between a surface- and a bulk term sets a critical radius above which a lumen can grow. In many biological systems, active processes contribute to these terms, i.e., cytoskeleton-generated cellular surface tensions and active pumping of fluid by the cells.

In many physical systems, heterogeneous nucleation – where an external wall or an impurity provides an additional interface – dominates over homogeneous nucleation. In classical nucleation theory, this is explained as a lowering of the free energy barrier that needs to be overcome for nucleation when an additional interface lowers the surface energy of the forming droplet (Turnbull, 1950). Here we investigate the role of an additional tissue in lumen formation. The external tissue acts like a wall on which heterogeneous nucleation can occur. We show how the presence of this additional interface facilitates lumen formation.

##### Critical Radius for Lumen Formation

We begin by deriving the critical radius above which a lumen will expand and below which it will disappear. This radius depends on the parameters of the system, i.e., the lumen pressure and the surface tension associated with the tissue-lumen interface. For a spherical lumen, the balance of forces at each point of the tissue-lumen interface is given by the law of Laplace:

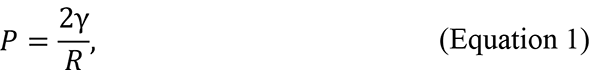

in which 𝑃 is the pressure difference across the tissue-lumen interface, γ is the effective surface tension of the tissue-lumen interface, and 𝑅 is the radius of the lumen. We assume that the pressure equilibrates within the cavity. In general, 𝑃 has several contributions, including active pumping terms. We decompose the pressure into a constant term 𝑃_𝐋_ and a dissipative term, and write the following constitutive relation:

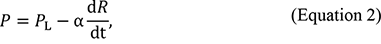

in which α is a dissipative coefficient associated with changes of the lumen radius. Here we do not consider any other dependencies of the pressure on the radius. From Equation 1 and Equation 2, we obtain a differential equation for the lumen radius:

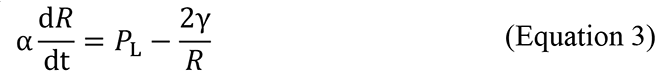

that has the traditional form of a nucleation equation with bulk and surface terms.

The critical radius for lumen growth is given by:

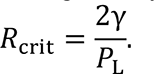

At which size a lumen will grow or shrink depends on the parameters γ and 𝑃_𝐋_ which are actively regulated by the cells through the formation of their apical domains and their pumping activities. In order to form a lumen, the bulk term must dominate over the surface term, i.e., active fluid pumping must overcome the cost of increasing the lumen-cell interface.

##### Presence of a Wall

We assume that the system is able to produce initial proto-lumina with a finite initialization volume 𝑉 - set for example by a characteristic exocytosis volume. The initial proto-lumen radius for the homogeneous case – i.e., in the absence of a wall – is then given by:

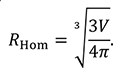

In classical heterogeneous nucleation, additional interfaces can lower the energetic cost of forming a new droplet interface, thereby reducing the work for nucleating the new phase at the boundary. A droplet in contact with a wall takes the shape of a spherical cap, for which the volume can be expressed in terms of the radius of curvature 𝑅_+.*_ and the contact angle 𝜃 (Figure 7F):

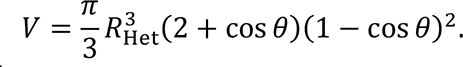

The contact angle is modified by the additional interfacial tensions with the wall as given by Young’s equation:

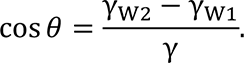

In the following, we denote by γ_1_ = γ_10_ − γ_12_, the difference between the surface tensions associated with the wall-tissue and wall-lumen interfaces, and we consider the regime of partial wetting, where γ_1_ < γ. The radius of curvature – which sets the surface term in Equation 3 – is thus given by:

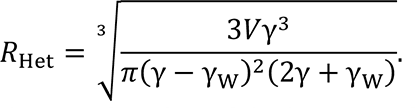

Figure 7G shows how 𝑅_+.*_ varies as a function of the lumen surface tension. Note that 𝑅_+.*_ > 𝑅_+,-_. The regimes of growth and shrinkage for the homogeneous and heterogeneous cases are depicted in Figure 7H.

In conclusion, if the interfacial tensions favor the formation of a new interface between the lumen and the wall, lumen expansion is facilitated by the presence of the wall.

### QUANTIFICATION AND STATISTICAL ANALYSIS

#### Image Analysis

Dimension measurements (Figures 3B, 3D and 5E), cell counts (Figures 3C, 3E, 5F, 7B, 7I, S1B-D, S2B and S2C), cell coordinates analysis (Figures 4A-C and S3A) and apical domain detection (Figure 6E) were performed with Imaris v9.2.1 (Bitplane). Signal intensity measurements (Figures 1D, 7C and 7D) and cell shape analysis (Figures 1B, 1G-I) were performed with Fiji (Schindelin et al., 2012). Cell tracking (Figure S3A) was performed with Fiji and Imaris. Whole embryo images were used for E4.5—E5.25 embryos. To compensate for signal attenuation in thicker samples, the half of the egg cylinder closer to the objective were used for dimension measurements and cell counts.

##### Evaluation of Embryo Morphology

Dimension measurements of *in utero* and 3D-geec embryos were performed in 3D using Measurement Points. For blastocysts, the diameter is defined as the mean of the long and short transverse axes of the ICM, and the length is defined as the distance between the epiblast-pTE boundary and the surface of the PrE. For egg cylinders, the diameter is defined as the mean of the long and short transverse axes of the egg cylinder, and the length is defined as the distance between the epiblast-extraembryonic ectoderm (ExE) boundary and the distal tip of the egg cylinder.

##### Evaluation of Embryo Development Based on Cell Numbers

Cell counts of *in utero* and 3D-geec embryos were performed in 3D using automated Spots detection with manual correction. Initial cell number (Figures S1B and S1C) was based on all nuclei stained by Hoechst 33342. VE (Figures 3C and 5F; also Figure 4) is defined as the visceral endoderm overlying epiblast cells; visceral endoderm overlying ExE is excluded from this analysis. Total cell number is defined as the sum of VE and epiblast.

To calculate the corresponding “*in utero* age” of D1 and D2 embryos based on their total cell numbers, a linear regression line was generated from the total cell numbers of *in utero* embryos (*y* = 75.06*e*^0.0712*x*^, R² = 0.933, where *y* is the total cell number (Figure 3E) and *x* is the age of the embryo in hours from conception; Figure S2B). Similarly, to calculate the normalized embryonic stage of live-imaged embryos (Figure 6C), a linear regression line was generated from the epiblast cell numbers of *in utero* embryos (*y* = 36.77*e*^0.0818*x*^, R² = 0.929, where *y* is the epiblast cell number (Figure 3C) and *x* is the age of the embryo in hours from conception; Figure S2C).

##### Evaluation of Cell Differentiation and Embryo Patterning at D2

VE cells were annotated for AVE identity by co-immunostaining of Gata4 and AVE markers, Lefty1 or Cerberus1, and their positions marked using automated Spots detection with manual correction. The distal tip of the egg cylinder was marked manually as a Spots object. A Reference Frame (X, Y, Z) was created with the *Z*-axis aligned along the proximal-distal axis of the egg cylinder. The 3D coordinates of the Spots in the given Reference Frame were used to calculate linear distance and angle of rotation about the *Z*-axis of each VE cell. E5.25 and E6.0 embryos were manually classified as Asymmetric, Symmetric or Borderline by qualitative AVE distribution. Polar plots of the spread of VE cells in each E5.25, E6.0 or D2 embryo were generated, with the distal tip as the origin, and converted to Cartesian coordinates. The centroid of the AVE cells for each embryo was calculated using the Cartesian coordinates. The distance of the centroid from the origin, scaled to the furthest VE linear distance of that embryo, was taken as the AVE Asymmetry Index of the embryo, with a value ranging from 0 to 1. As all Symmetric embryos have an AVE Asymmetry Index lower than 0.15, this value was taken as the threshold for evaluation of 3D-geec embryo development at D2 based on AVE asymmetry. The AVE Asymmetry Index for each D2 embryo was calculated as described, and the percentage of embryos evaluated as successfully body axis specified was 67% (*n* = 18, *N* = 6) (Figure 4).

##### Dynamics of Pro-amniotic Cavity Formation

Apical domain detection was performed in 3D using manual Spots generation. Spots objects were generated based on Ezrin-mCherry reporter signal, and their 3D coordinates were used to calculate their distance from the embryo center, scaled by the average radius of the epiblast tissue at the respective time points (Figure 6E).

##### Analysis of Cell Dynamics in the Epiblast

Ten epiblast cells in a cluster before ExE invagination in an H2B-GFP;mT embryo were tracked over 16 hours. The mean displacements of these cells in 3D from the centroid over time were calculated as an indicator of cell dispersion (Figure S3A).

##### Analysis of Signaling Activity in the Epiblast

Immunostaining of Id1 was used as a readout for BMP signaling. The reporter lines Dusp4- T2A-mVenus and A7-Venus were used as readouts for FGF-Dusp4 and Nodal-Foxh1 signaling, respectively. As signaling activity is heterogeneous within the epiblast tissue, we used a proportion of cells in a high expression state as a readout of local signaling activity in the tissue. Intensity measurements were performed on a Z-projection of 10 µm slices around the equatorial plane of the embryo, processed to subtract background using a rolling ball of radius 100 px (pixel size of 0.165 µm/px). Circular ROIs were drawn on epiblast nuclei in the resultant image so that each ROI was of the largest diameter that fits in the nucleus, and care was taken to avoid including regions where two nuclei overlapped due to Z-projection. The mean signal intensity in the Id1 or Dusp4-T2A-mVenus and Hoechst channels of each ROI was measured, along with the coordinates of its center. ROIs were segregated into either the proximal half or the distal half of the epiblast based on their position. Due to epiblasts being misshapen in the absence of ExE, the left and right 20% of cells were discarded from the analysis. A cell with a Hoechst-normalized Id1 or Dusp4-T2A-mVenus signal intensity higher than the median value of all epiblast cells in the embryo was annotated as Id1^high^ or Dusp4^high^. The percentage of Id1^high^ or Dusp4^high^ cells in the proximal and distal halves of each embryo was calculated and correlated with each other by a solid black line to represent the signaling landscape (Figures 7C and 7D).

#### Machine-learning-based Segmentation and Analysis

The segmentation pipeline used to process the 3D images of the mTmG membrane signal consists of four steps. In the first step the 3D input images are pre-processed, where every Z-slice is down-sampled by a factor of 4 along each axis by locally averaging squares of 4×4 pixels. The resulting images display the same physical volume with a dimension of 512×512×400 voxels and a physical voxel size of 0.832×0.832×1.000 µm^3^ (X,Y,Z). In the second step a neural network implementation from PlantSeg is used to generate a probability map of the membrane locations. In the third step the probability maps are segmented using a set of algorithms provided by PlantSeg (Wolny et al., 2020). The best segmentation algorithm and its corresponding hyper-parameters were found by a custom-made pipeline which explored thousands of different configuration parameters simultaneously using EMBL’s computing cluster. In the fourth step the best segmentation output is manually selected through visual inspection and used for further analyses.

Since no ground truth segmentation was initially available, performance of the complete segmentation pipeline was improved by the following iterative procedure. In the first iteration a pre-trained neural network available in the PlantSeg package was used to generate the initial membrane probability maps. In particular, we used a CNN trained on the Arabidopsis ovules dataset (https://osf.io/w38uf) named “confocal_unet_bce_dice_ds2x”.

Having the cell boundary prediction, the initial segmentation was produced with PlantSeg. The segmentation results were improved by choosing the most correctly segmented volumes (inspected visually) and using them as ground truth labels to train a dedicated neural network for the membrane prediction task. The process of choosing the best segmentation results and re-training the network was performed twice.

Analysis of cell parameters was performed using Python 3.8 based on the segmentation generated by the above process. From five representative H2B-GFP;mT embryos live-imaged (Figure 5B), 10 well-segmented cells per embryo were picked up per hour until pro-amniotic cavity expansion (Figure 6B) and used for analysis with care taken to avoid bias in the cells sampled (Figure 6C). Aspect ratio is calculated by fitting an ellipsoid to the cell and dividing the longest axis (LA) of the ellipsoid by its shortest axis (SA). Long-axis alignment is calculated as the angle between the LA of the cell and a line segment connecting the outermost voxel of the cell to the center of the cell. A low angle indicates an alignment of the long axis along the inside-outside axis of the egg cylinder.

#### Derivation of Optimal Initial Cell Number Threshold

A confusion matrix for each threshold level from 0 to 230 cells (with intervals of 10 cells) was constructed with the following definitions:

True positives (TP): D0 embryos *above* a threshold that yield egg cylinders

False positives (FP): D0 embryos *above* a threshold that do *not* yield egg cylinders True negatives (TN): D0 embryos *below* a threshold that do *not* yield egg cylinders False negatives (FN): D0 embryos *below* a threshold that yield egg cylinders.

The threshold level that yields the highest Accuracy (i.e., (TP + TN)/(TP + FP + TN + FN)) provides the best tradeoff between sample retention and egg cylinder formation efficiency. As such, a threshold of ≥110 cells provided the highest optimality, qualifying 66% of E4.5 embryos recovered from natural mating and resulting in 74% egg cylinder formation efficiency (*n* = 35, *N* = 4) (Figure S1C).

#### Statistical Analysis and Data Reproducibility

Experiments in this study were performed at least at three independent times, except for the data shown in Figures S1D, S3A and S3B. *N* values represent the number of independent experiments, while *n* values represent the total number of embryos collected from independent experiments. Performance of 3D-geec was independently replicated by two operators, T.I. and HT.Z. Data analysis and statistical tests were performed in Rstudio or

OriginPro. Details of the statistical analysis are provided in the figure legends. Briefly, the normality of the distribution for each dataset was tested by the Shapiro-Wilk test. When the data followed a normal distribution, difference among groups in comparison was examined by either *t*-test (for comparison of two groups) or one-way ANOVA (for comparison of more than two groups) followed by Tukey’s post hoc test. Otherwise, nonparametric Kruskal-Wallis ANOVA was used with Mann-Whitney *U*-test. No statistical method was used to predetermine the sample size. Experiments were not randomized, and the investigators were not blinded to allocation during the experiments and outcome assessment. Data visualization was also performed in Rstudio using the ggplot2 package or OriginPro. Box and whisker plots show the following: boxes represent the 25^th^ and 75^th^ percentile range, whiskers represent the 1.5x interquartile range.

#### Analytical Calculations

Analysis and plotting for Figures 7F-H was performed with Wolfram Mathematica 12.1.1.0.

## Supplemental Tables

**Table S1.**
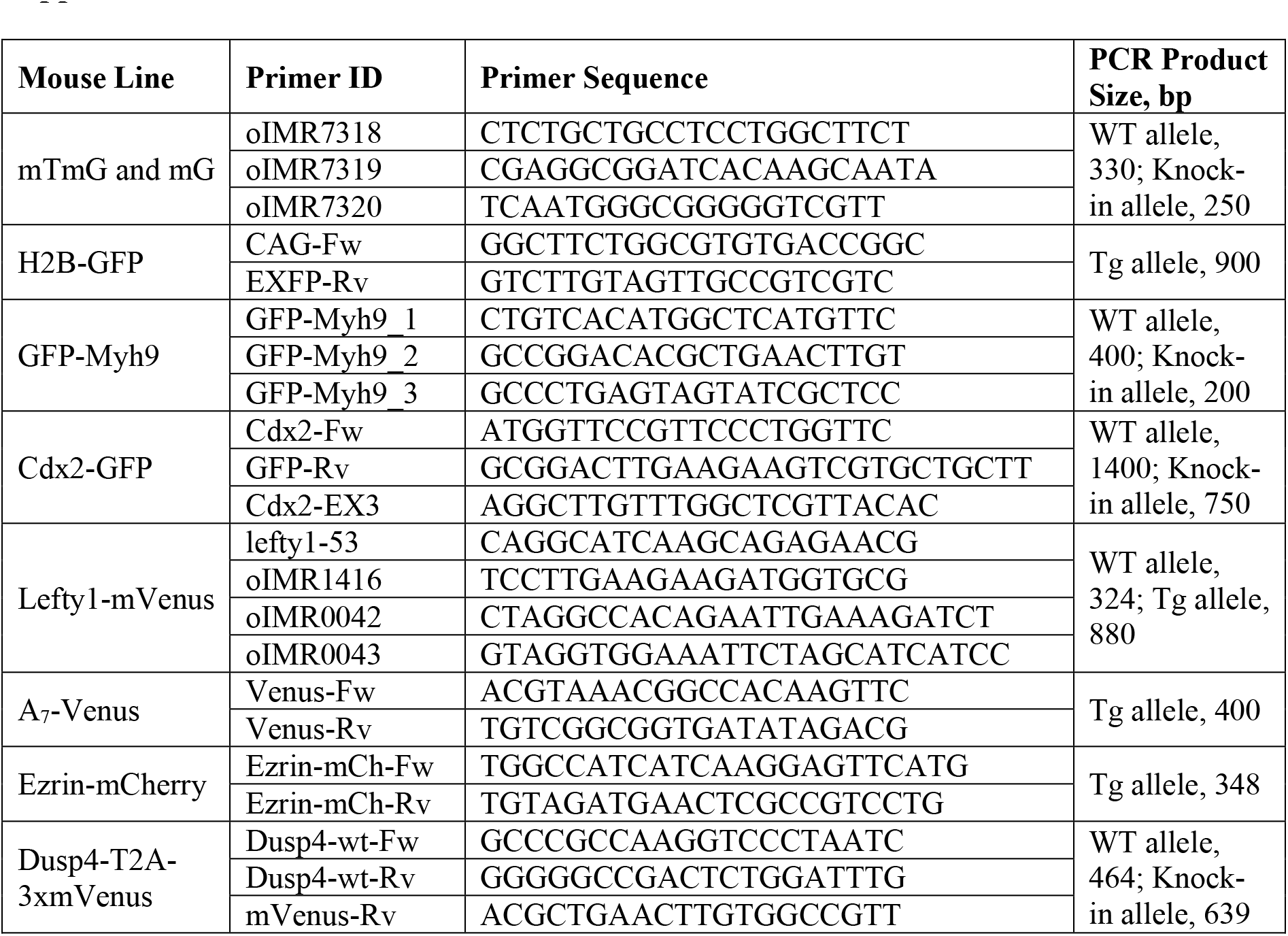
Genotyping primers and PCR product sizes, Related to STAR Methods.

## Supplemental Videos

**Video S1. Laser ablation induces apical constriction of pTE cells over the following 20 minutes, Ralated to** Figure 1.

Time-lapse images of a GFP-Myh9 embryo after 6 hours of 3D-gel culture, ablated with infra-red laser pulses. Ablated point is marked with an asterisk (*t* = 00:00). Gray, GFP-Myh9. Time, minutes:seconds. Scale bar, 20 µm

**Video S2. Microsurgery excising the mTE induces invagination of pTE cells after 2 hours, Ralated to** Figure 1.

Time-lapse images of a Cdx2-GFP embryo at E4.5 following microsurgical removal of mTE cells. Green, Cdx2-GFP. Time, hours:minutes:seconds. Scale bar, 20 µm

**Video S3. Light-sheet live-microscopy reveals 3D-geec embryos develop into the egg cylinder, Ralated to** Figure 5.

Time-lapse images of an H2B-GFP;mT embryo developing in 3D-geec from D0 to D2. Note that the equatorial plane of the embryo at each time point is shown in Figure 5B. Green, H2B-GFP; magenta, mT. Time, hours:minutes. Scale bar, 50 µm

**Video S4. 3D-geec recapitulates the anterior-posterior axis by migration of Lefty1^+^ distal visceral endoderm cells to the anterior,**

**Ralated to Figure 5.**

Time-lapse images of a Lefty1-mVenus;mT embryo developing in 3D-geec during D1-2. 3D rendering by Imaris. Green, Lefty1-mVenus; gray, mT. Time, hours:minutes. Scale bar, 50 µm

**Video S5. Machine-learning-based cell membrane segmentation allows for the analysis of cellular dynamics,**

**Ralated to** Figure 6.

Time-lapse images of an H2B-GFP;mT embryo developing in 3D-geec during the first 24 hours (left), its output of automatic cell membrane segmentation (middle) and a merged view of the membrane (gray) and segmentation (Glasbey) (right). Time, hours:minutes. Scale bar, 50 µm

**Video S6. The apical domain emerges in epiblast cells and progressively clusters at the center of the epiblast tissue,**

**Ralated to** Figure 6.

Time-lapse images of an Ezrin-mCherry;mG embryo developing in 3D-geec from 6 to 28 hours until pro-amniotic cavity formation. Green, Ezrin-mCherry; magenta, mG. Time, hours:minutes. Scale bar, 50 µm

---

**Figure S1.**
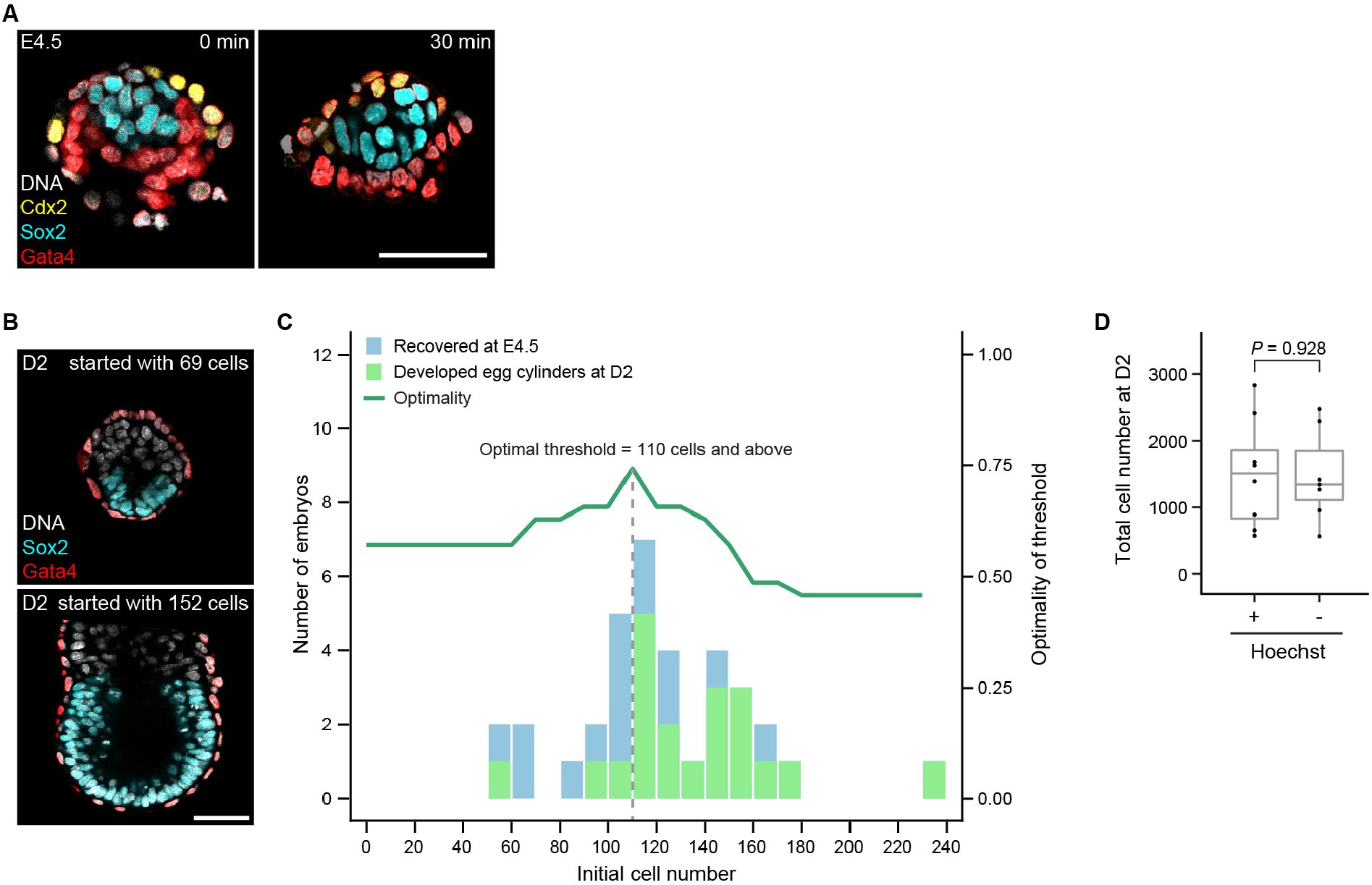
Quality control for E4.5 mouse blastocysts suitable for 3D-geec, Related to Figure 2, 3 and 4. (A) Immunofluorescence of representative E4.5 blastocysts upon mechanical dissection of mTE (left) and after 30 minutes incubation in IVC1 medium (right) stained for Sox2^+^ epiblast, Gata4^+^ VE and Cdx2^+^ TE. Note the tissue shape change, due possibly to pTE contraction. (B) Immunofluorescence of 3D-geec embryos at D2 with initial cell number 69 (top) and 152 (bottom) stained for Sox2^+^ epiblast and Gata4^+^ VE. (C) Histograms of the number of embryos recovered at E4.5 (blue) and those developed to egg cylinder by 3D-geec at D2 (green) for a given total number of ICM and pTE cells (bin size = 10), with a line chart of optimality (see STAR Methods). A threshold level of ≥110 cells provides the best optimality of 0.74. *n* = 35; *N* = 4. (D) The total epiblast and VE cell number of Hoechst-treated and -untreated 3D-geec embryos at D2. Hoechst-treated: *n* = 8, Hoechst-untreated: *n* = 7; *N* = 2. *P* value calculated using Mann-Whitney *U* test. Scale bars, 50 µm.

**Figure S2.**
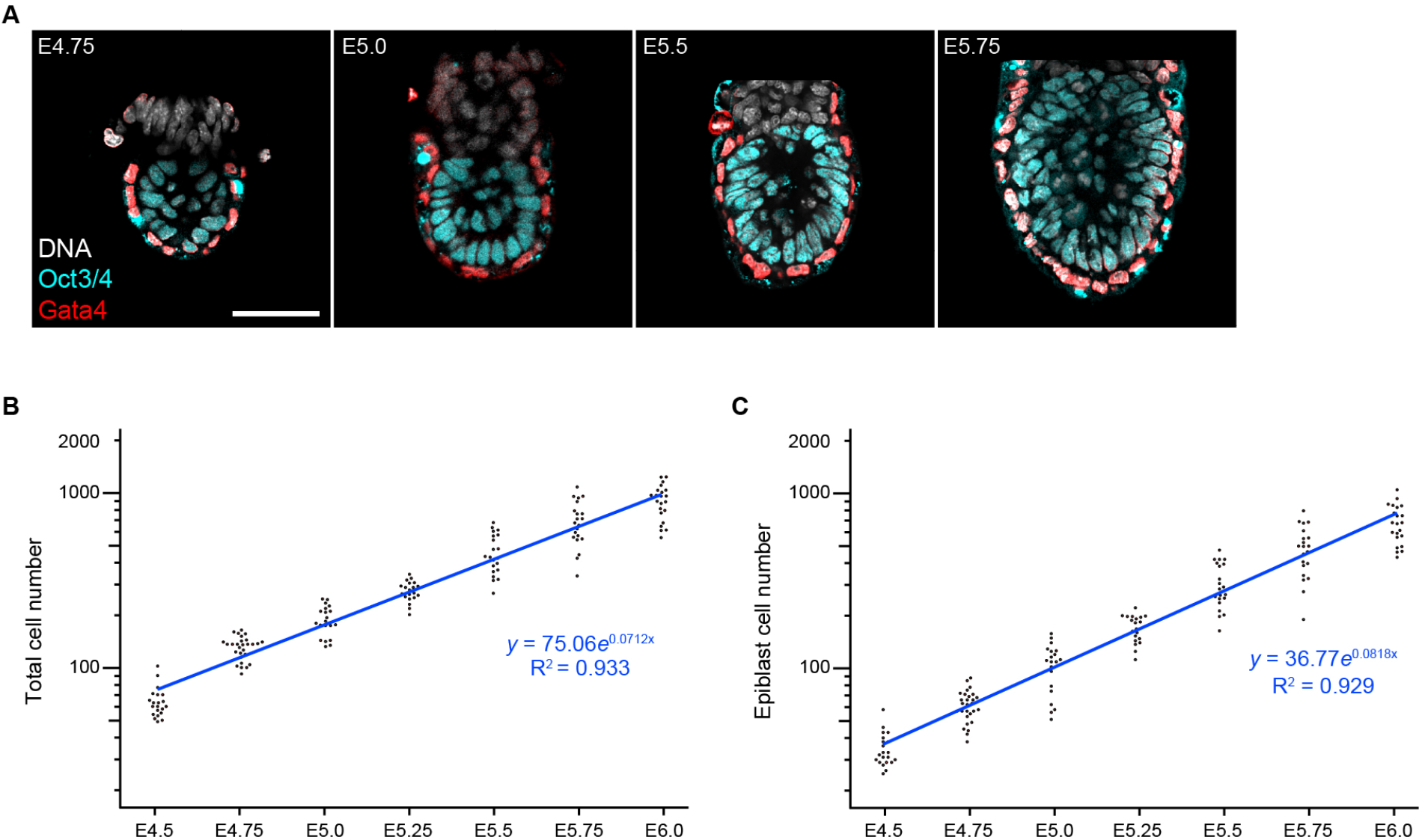
Quantitative characterization of peri-implantation mouse embryos, Related to Figure 3 and 6. (A) Immunofluorescence of a representative embryo developed *in utero* at E4.75, E5.0, E5.5 and E5.75 stained for Oct3/4^+^ epiblast and Gata4^+^ VE. (B, C) Plots showing the total (epiblast and PrE/VE) cell number (B) or epiblast cell number (C) against the developmental stage of the recovered embryo defined by the time of recovery. Based on the cell numbers and the regression lines, embryos can be re-defined as a quantitatively normalized stage (nE; see Figure 6C and STAR Methods). Equation of regression line for total cell number is *y* = 75.061*e*^0.0712*x*^; that for epiblast cell number is *y* = 36.77*e*^0.0818*x*^. *n* = 21 (E4.5), 28 (E4.75), 20 (E5.0), 20 (E5.25), 21 (E5.5), 21 (E5.75) and 22 (E6.0). Scale bar, 50 µm.

**Figure S3.**
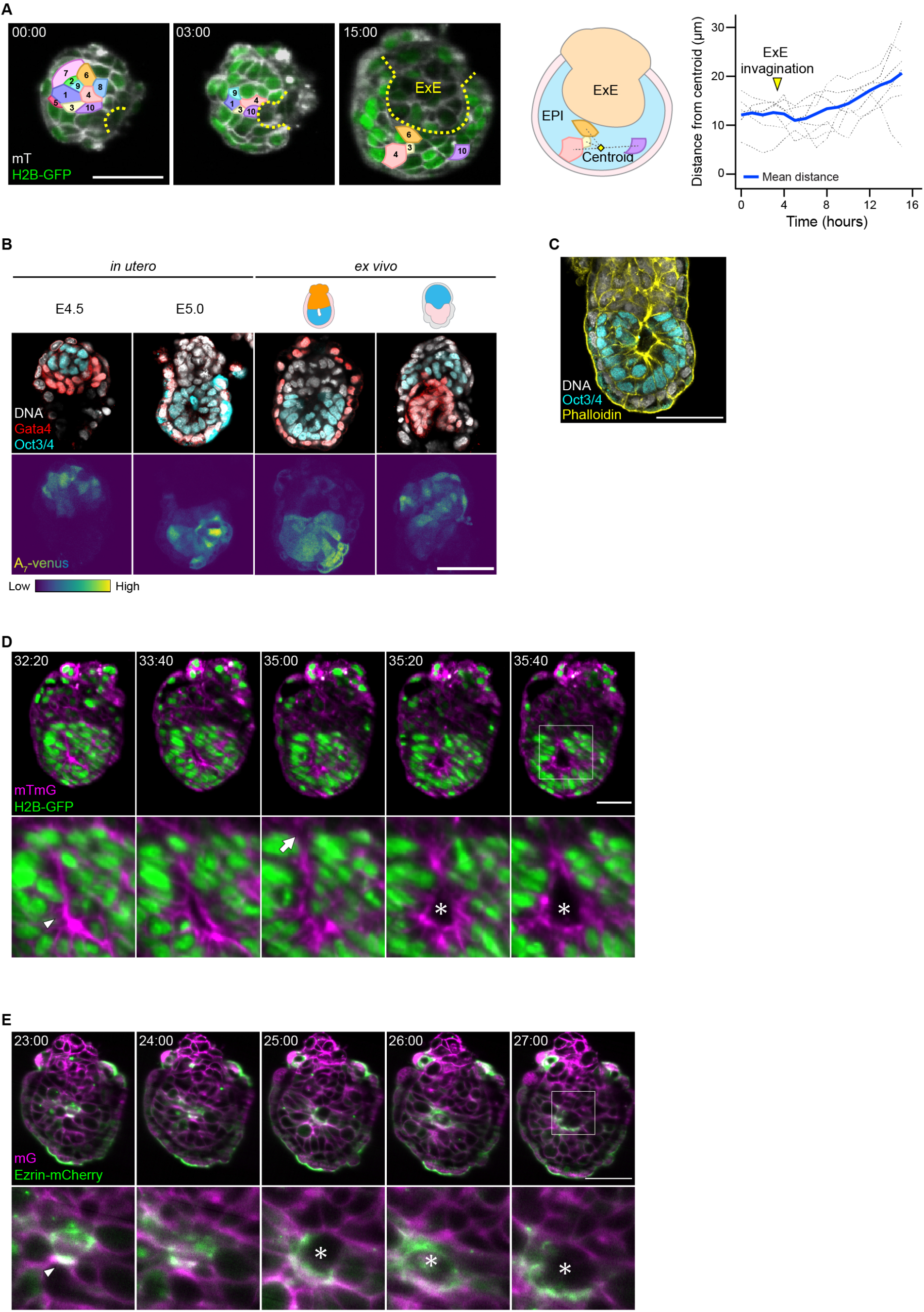
Impact of ExE on epiblast cell dynamics and Nodal signaling, Related to Figure 7. (A) Tracking of 10 neighboring epiblast cells in a representative H2B-GFP;mT embryo during the first 15 hours of 3D-geec culture. Mean distance from the centroid of tracked cells is used as a measure of cell dispersion. *t* = hours:minutes. Yellow broken lines mark the boundary between ExE and epiblast. (B) Immunofluorescence of representative A7-Venus embryos developed *in utero*, or *ex vivo* in the presence and absence of ExE, stained for Oct3/4^+^ epiblast and Gata4^+^ VE. *n* = 3, 8, 2 and 3, respectively. (C) Immunofluorescence of a representative E5.0 embryo developed *in utero*, stained for actin and Oct3/4^+^ epiblast. (D, E) Time-lapse images of representative H2B-GFP;mT (D) and mG;Ezrin-mCherry (E) mouse embryos developed *ex vivo* in the presence of ExE from E4.5 (*t* = 00:00, hours:minutes). Arrowheads and asterisks mark rosettes and the pro-amniotic cavity, respectively. An arrow indicates a fissure-like structure connecting to the boundary between epiblast and ExE. Scale bars, 50 µm.

